# The importance of mistakes: Variation in spores reveals trade-offs for range expansion in the xeric-adapted Australasian species *Cheilanthes distans* (Pteridaceae)

**DOI:** 10.1101/2024.06.22.600204

**Authors:** Karla Sosa

**Affiliations:** New York University

## Abstract

Biological trade-offs present a central issue for evolutionary biology: it has been a fundamental understanding within the field that limits exist on the phenotypic traits a species is able to exhibit in part due to trade-offs. Reproduction—with its myriad forms—has been studied extensively in the context of these dynamics. And while considerable literature has explored trade-offs between seed size and number and their associated environmental conditions, none has looked at spore size trade-offs in ferns. We can hypothesise potential trade-offs in spore size: smaller spores should be able to disperse farther, but may not have sufficient provisions to survive in environments that require them to remain at the gametophyte stage for longer periods if their germination cues are mismatched. Reproductive mode (sexual vs. asexual) and ploidy may also be playing a role. In order to study trade-offs related to spore size, I focus on the Australasian fern species *Cheilanthes distans* (Pteridaceae), which is most often found in xeric environments, growing in crevices or on top of rocks which are haphazardly scattered across their range. Apomictic diplospores in this species are formed through first division restitution, a meiotic pathway particularly prone to mistakes in chromosome pairing and cell division (as compared to premeiotic endomitosis). Rather than being problematic, these mistakes ultimately lead to considerable additional variation in spore size, spore products (through a range of aneuploid spores), and spore ploidy. In this study, I explore trade-offs between spore size, dispersal, and germination, taking into account effects from reproductive mode and ploidy. I carried out an extensive survey of *C. distans* specimens to establish the prevalence of sexual vs. apomictic (asexual) specimens, and to describe in greater depth the variation in ploidy across the species. I also collected data on spore size and sporogenesis forms. With these data I then asked: is spore size correlated with range area or with germination? And does spore form correlate with either spore size or germination? Ultimately, I find that variations in sporogenesis may be leading to large variation in spore sizes—especially since spores traditionally considered abortive are in fact viable—and that this variation may provide abundant fodder for evolution to act through trade-offs between dispersal into large ranges and germination leading to establishment. Especially in light of the fact that many spores that were historically considered abortive are fully viable and likely shaping evolution in important ways, it is worth remarking on what these results illustrate more broadly: the way in which we have constructed ‘disability’ ultimately affects how we perceive so-called ‘genetic errors’—both in humans and in other species—and thus limits what we allow ourselves to imagine ‘disabled’ beings are capable of.

## 1. Introduction

Biological trade-offs present a central issue for evolutionary biology (Stearns 1989; Roff and Fairbairn 2007 and references therein). In its most essential form, a trade-off occurs when an increase in one trait causes a decrease in a different trait; their respective increases and decreases may or may not be linked with Darwinian fitness (Garland 2014), although those that do affect fitness are of greatest interest to evolutionary biologists. Among these, there are two main types: phenotypic trade-offs and genetic trade-offs, the latter involving negative covariances between genetic elements (Roff and Fairbairn 2007). If we focus our attention on phenotypic trade-offs, fundamental understanding within evolutionary biology is that limits exist on the traits a species is able to exhibit in part due to trade-offs. A classic mechanism that captures these limits is the ‘Y-model’, which states that a particular unit of resource (e.g. time, energy) can only be used towards a single trait at a time (e.g. an hour spent foraging cannot be spent building a nest or finding a mate) (the Y-model’s historical use and derivation is explained in Roff and Fairbairn 2007). Given how fundamental these limits are, their study can both elucidate constraints experienced by organisms as well as point to aspects of a species’ biology that are under strong evolutionary pressure.

Studying trade-offs is important as we seek to understand variation in a species. While it is true that trade-offs aren’t per se causing variation—which as we know is a product of genetic and environmental effects—it is precisely the existence of environmental variation that may lead different states of a trait (i.e. smaller or larger, fewer or more, etc.) to be favoured at different times and places (Stearns 1989), thus leading us to observe it in nature. However, the variation—and in particular the combinations of variation across different traits—we observe may be limited by trade-offs. While it may be possible for an organism to physically achieve a particular state (say, very large or very small), if opposing trade-offs exist between this and a different trait (say number), any given state may not be achievable in combination with a subset of states in the other trait (say very large can only exist with few, or very small with many). While it is possible for this dynamic to be disrupted—in particular by novel mutations that release constraints (e.g. Flagel and Wendel 2009)—trade-offs will nevertheless play an important role in determining the combinations of traits we observe in a species.

Reproduction—with its myriad forms—has been studied extensively in the context of trade-offs. One of its central theories is r/K selection (Pianka 1970), which posits trade-offs between number of offspring and parental investment, sometimes shorthanded as quality of offspring. The balance of where a species lands in this trade-off will depend on their specific biology and which strategies confer the highest fitness: for example, organisms in unpredictable environments will often decrease offspring number but increase parental investment (Husby 1986; Badyaev and Ghalambor 2001), while organisms from ephemeral environments or those frequently disturbed tend to produce many offspring with relatively lower investment per offspring (Eckerström-Liedholm et al. 2017; Morrongiello et al. 2012). In the case of plants, parental investment is often measured via the size of propagules (i.e. seeds and spores). Considerable literature has explored trade-offs between seed size and number, and the associated environmental conditions leading to particular trade-off dynamics (e.g. Venable 1992; Leishman 2001; Sadras 2007; Muller-Landau 2010). However, only a single study has looked at spore size trade-offs in plants (Löbel and Rydin 2010, which studied epiphytic bryophytes), and none has assessed spore size trade-offs in ferns, which merit study separate from bryophytes given their extremely different reproduction strategies. Bryophytes are in some ways more similar to seed plants since sexual reproduction occurs embedded within the dominant generation (Longton 2019), such that—for most species at least—each plant can provide greater parental care (be it through resources or protection) during this process. By contrast, sexual reproduction in ferns occurs in free-living gametophytes that are very small and delicate (Moran 2004), and they must survive and achieve reproduction with no parental investment beyond the contents packed within a spore.

The dearth of studies on ferns may be due to the half-lore idea that fern spores can be dispersed freely by the wind and find their way to any available habitat space (e.g. Tryon 1970). Presence of a particular species is then solely determined by habitat. To quote Laurens Baas Becking on microorganisms: “*Everything is everywhere, but, the environment selects*” (de Wit and Bouvier 2006). Among vascular plants, the propagules of ferns are in fact particularly dispersible given the small sizes of spores and their light weight, which is lighter than the smallest seeds and even than the pollen of several wind-dispersed species (Gómez-Noguez et al. 2016 and references therein). It has been theorised that spores can be carried essentially anywhere on the planet (Tryon 1970; Wolf et al. 2001); their higher proportions relative to other vascular plants on oceanic islands (vs. the mainland) in part supports this conclusion (Tryon 1970; Smith 1972; Peck et al. 1990). If we assume that everything is in fact everywhere with relation to fern spores, then spore variation becomes unimportant—and thus unworthy of study—because we are assuming that spores of all sizes will arrive at any given location. While we should not discount the importance of extremely rare events in shaping history, we should not assume that simply because it seems theoretically possible for fern spores to arrive at all available habitat space, that they will in fact do so.

The predictions in the papers cited above have not, for example, considered that larger spores are expected to settle closer to their source, as explained by Stokes’ law (Stokes 1880). Empirical studies have found this to be true for both spores and seeds (Norros et al. 2014; e.g. Muller-Landau et al. 2008; Wilkinson et al. 2012 and citations therein). We know also that variation in spore size exists in ferns, with larger spores often having higher ploidy levels (e.g. Sigel et al. 2011; Huang et al. 2006; Beck et al. 2010). Further, ferns exhibit variation in the way spores are produced, including modifications to typical meiosis that result in the formation of “diplospores” (spores that are diploid rather than haploid), which germinate into apomictic organisms. Altered spore formation often leads to *additional* variation in spore size due to frequent mistakes during modified meiosis that result in variously aneuploid and abortive spores (Braithwaite 1964; Walker 1985; Grusz 2016; Manton 1950). Given this, we should expect to see variation in the distances that fern spores are regularly dispersing.

We must also consider the possibility that spore size may be under selection. In fact, Carlquist posited that fern spores were unusually large in certain taxa (Schizaeoideae, Hymenophylloideae, Adiantoideae, Blechnoideae) found on Hawaiian islands so as to reduce their dispersal into unsuitable environments or out to sea (Carlquist 1966). Even so, we should still expect dispersal to be broadly favoured, as high densities of offspring near maternal plants affect survival and growth of offspring through competition, predation, and pathogens (Augspurger 1983; Clark and Clark 1984; Platt 1976). We might also expect longer distance dispersal—which may lead to range expansion^1^—to be advantageous in cases where suitable habitat is narrow or ephemeral, and scattered across the landscape (Johst et al. 2002; Saura et al. 2014), such as is the case in epiphytes, lithophytes, or serpentine plants (Löbel and Rydin 2009; Spasojevic et al. 2014). As a matter of fact, we find that many fern species are widely distributed (Smith 1972), such that frequent dispersal must have occurred at least historically.

For plants, successful reproduction—whether or not it involves range expansion—involves two key steps: dispersal and establishment. Spore size is likely to affect dispersal. Size is also likely to affect establishment because larger spores should provide greater resources for germinating gametophytes, which must survive independently from the parental plant and develop sufficiently in order for a successful embryo to develop (Gemmrich 1977; Ballesteros and Walters 2007). The resources packed into the spore may thus be especially important for gametophyte growth and survival depending upon the environment, since harsher environments may prove more difficult for proper resource acquisition for such small, typically short-lived organisms (Moran 2004). Further, spore provisioning may also be important depending upon the type of reproductive strategy a gametophyte will undertake. Ferns—sometimes even those within the same species—have an array of reproductive strategies available to them (Haufler et al. 2016). One of these is outcrossing, which occurs between two gametophytes from different parental sporophytes. This strategy will preserve the greatest genetic variation in a population, so it is generally favoured and the strategy most frequently observed in natural populations (Sessa et al. 2016). A second strategy is sporophytic selfing, which occurs between two gametophytes from the same parental sporophyte, and is akin to selfing in flowering plants. For reproduction to happen successfully for both outcrossing and sporophytic selfing, there needs to be a continuous film of water between both gametophytes so that the sperm from one gametophyte can swim to the egg from a separate gametophyte. A third strategy is gametophytic selfing, where a single gametophyte fertilises itself, leading to a completely homozygous embryo but requiring only a small film of water for the sperm to travel in. A final strategy is apomixis, or asexual reproduction, which does not allow for genetic recombination but does not require additional water beyond that needed for baseline gametophyte survival. Spore provisioning may thus be particularly important in low-water conditions and for sexually-reproducing gametophytes, as these must grow large enough to survive until sufficient water is available for final development and reproduction (Moran 2004). We can thus hypothesise a potential trade-off in spore size: smaller spores should be able to disperse farther, but may not have sufficient provisions to survive in unfamiliar environments that may require them to remain at the gametophyte stage for longer periods if their germination cues are mismatched to this new environment. Further, reproductive mode (sexual vs. asexual) and ploidy (given that higher ploidy levels have larger spores) may also be playing a role when it comes to trade-off dynamics related to spore size.

In order to determine whether this type of trade-offs occurs, I have chosen to focus on the Australasian fern species *Cheilanthes distans* (Pteridaceae). This species presents an ideal system given its widespread distribution across Australia as well being found in New Zealand/Aotearoa (including in the Kermadec Ecological Region), New Caledonia, Lord Howe Island, Norfolk Island, and Papua New Guinea (Tindale and Roy 2002). *Cheilanthes distans* is most often found in xeric environments, growing in crevices or on top of rocks which are irregularly scattered across their range (Quirk et al. 1983; Sosa, pers. obvs.). These characteristics should allow us to test whether spore provisioning may be of importance for offspring survival for organisms living in harsh environments. Of the 21 specimens that had been previously examined, all were found to be apomictic, and cytological work had found *C. distans* to be a triploid except for an uncertain tetraploid individual, nevertheless hinting at potential cytological variation in this species (Quirk et al. 1983; Chambers and Farrant 1991; Tindale and Roy 2002). It is worth noting that apomictic diplospores in this species are formed through first division restitution (Tindale and Roy 2002). This pathway has been noted as being particularly prone to mistakes in chromosome pairing and cell division during meiosis, especially as compared to premeiotic endomitosis (Braithwaite 1964; Walker 1985; Mehra and Singh 1957; Hickok and Klekowski Jr. 1973; Grusz 2016). Rather than being problematic, these mistakes—which I will refer to as spore forms^2^—ultimately lead to considerable *additional* variation in spore size, spore products (through a range of aneuploid spores), and possibly spore ploidy.

For my study, I sought to explore trade-offs between spore size, dispersal, and germination, taking into account effects from reproductive mode and ploidy. For this I carried out an extensive survey of *C. distans* specimens to establish the prevalence of sexual vs. apomictic (asexual) specimens, and to describe in greater depth the variation in ploidy across the species. I also collected ample data on spore size and sporogenesis forms, which I will refer to as spore forms hereafter. With these data I then asked: is spore size correlated with range area or with germination? I also aimed to answer: does spore form correlate with either spore size or germination? Ultimately, I find that variations in sporogenesis may be leading to large variation in spore sizes—especially since spores traditionally considered abortive are in fact viable—and that this variation may provide abundant fodder for evolution to act through trade-offs between dispersal into large ranges and germination leading to establishment.

## 2. Materials and Methods

### 2.1 Specimens

Herbarium specimens for *C. distans* were requested on loan from herbaria with the most significant holdings for this species, namely: State Herbarium of South Australia (AD), Auckland War Memorial Museum (AK), Queensland Herbarium (BRI), Australian National Herbarium (CANB), Manaaki Whenua – Landcare Research (CHR), Australian Tropical Herbarium (CNS), Tasmanian Museum and Art Gallery (HO), Royal Botanic Gardens Victoria (MEL), University of New England (NE), Royal Botanic Gardens and Domain Trust (NSW), Western Australian Herbarium (PERTH), and Smithsonian Institution (US). Record information for most of these specimens was downloaded from Australia’s Virtual Herbarium database (Herbaria 2021) in October 2018. For US specimens, which are not included in this database, records were entered manually. For specimens that were not georeferenced I attempted to locate their coordinates based on label information and using GEOLocate (Rios 2021) and Google Maps. Existing georeferenced specimens were examined visually to remove any outliers or centroid values (i.e. values georeferenced at the level of a region, for example “Queensland”). Any specimens that could not be given coordinates were removed from the study. Apologies and amends are owed to the Traditional Custodians and Indigenous Nations from whose lands these samples and those below were taken without consent nor compensation.

These samples were complemented with pressed, oven-dried specimens collected in the field during February–March 2019. These samples were collected near Auckland, New Zealand/Aotearoa, and in the vicinity of Cairns and Queenstown, Queensland, Australia.

Because there were more specimens than I could examine, I focused my sampling on herbaria of particular interest (i.e. PERTH and BRI, as both southwestern Australia and Queensland are some of the wetter regions in Australia, where sexual specimens may thrive) and otherwise aimed to sample evenly across herbaria, across geography (this was achieved by visually checking sampling on a map), and across morphology.

### 2.2 Locating the sexual diploids and characterising spore variation

#### 2.2.1 Spore data collection

Spore number per sporangium was counted to determine reproductive mode. In *Cheilanthes s.s*., a specimen is presumed sexual if it contains 32 spores per sporangium, and apomictic if it contains 16 spores per sporangium (Tindale and Roy 2002; Sosa et al. 2021; Smith 1974). Intact sporangia were located on each specimen. Each intact sporangium was transferred with a dissecting needle to a drop of glycerol on a glass slide. The sporangium wall was then carefully ruptured, allowing the contents of the sporangium to be released within the glycerol and for an exact spore count to be determined. I aimed to sample three mature, intact sporangia per specimen. It was not always possible, especially as some sporangia looked intact but were slightly broken either prior to or upon transfer. These cases were recorded along with final counts.

The resulting spore preparations were photographed at 20x using an ECHO Revolve confocal microscope (Discover Echo Inc., USA). This microscope displays images on a screen rather than through eyepieces, and this screen and the built-in software can also be used to capture and annotate images. Spore diameters were measured directly on these images and recorded as annotations using the ECHO Revolve software. Only spores that appeared viable (for this, see criteria in 2.4) and had not been broken during dissection were measured. These measurements were then used to estimate total population as well as within sample variances in R version 3.6.0 (R Core Team 2017). Spore images were also used to assess spore forms (see 2.4).

All spore images and other associated data can be found at the following repository: https://doi.org/10.7910/DVN/NFI0LE. (I collected similar data for *Cheilanthes tenuifolia*, but ultimately did not use it. You can find those data at the following repository: https://doi.org/10.7910/DVN/S6BGHN – please use it!) The code used for the analyses below can mostly be located here: https://doi.org/10.7910/DVN/EPDSZN. The data and code used for Section 2.2.2 is instead located here: https://doi.org/10.7910/DVN/AUHWGP.

#### 2.2.2 Ploidy estimation

I attempted to grow plants from herbarium specimens to determine ploidy using chromosome squashes. Spores from herbarium specimens for which measurements had been obtained and that were collected in 1990 or later, as well as from all presumed sexual specimens, were sown under a flow hood onto petri plates with Hevly’s medium (Hevly 1963) and sealed with Parafilm. Plates were kept at room temperature with a 12h–12h light–dark cycle under 39 µmol s^-1^ m^-2^ per µA of fluorescent light, measured with a LI-COR LI-250A light meter (LI-COR Biosciences, Nebraska, USA). Young sporophytes were transferred to plastic pots with Biocomp BC-5 soil kept under the same light conditions and watered once a week. Chromosome counts were obtained using standard protocols for meiotic material (Windham and Yatskievych 2003) and results were documented using a digital camera (EOS Rebel T3i, Canon Inc., Japan) mounted on a phase contrast microscope (MT5310L, Meiji Techno Co., Japan). In order to respect and honour Indigenous peoples’ relationship and rights to their close non-human relatives, sporophytes from this and subsequent experiments were autoclaved after protocols were completed.

Because chromosome squash data collection was limited by the COVID-19 pandemic, I also estimated ploidy using nQuire (Weiß et al. 2018), which estimates ploidy based on sequence data and which has been shown to be accurate on herbarium specimens (Viruel et al. 2019). Sequence data for *C. distans* were primarily collected for the work outlined in Sosa, 2024, and greater detail on methods is provided there. Methods are outlined here briefly: DNA was extracted from herbarium specimens or, when available, from silica-dried leaf material collected in the field. DNA was extracted using Omega EZNA SP Plant DNA Kit (Omega Bio-tek, Georgia, USA) with standard protocols. Extractions were sequenced at RAPiD Genomics using GoFlag 451 target enrichment probes (Breinholt et al. 2021). Resulting sequences were then filtered using HybPiper (Johnson et al. 2016) for the analyses in Sosa, 2024. One of the outputs generated by HybPiper includes the number of paralogs for each of the loci. These results were used to select loci for use in the nQuire analyses; only loci with at most four paralogs were included in the ploidy estimation. The reasons for this cut-off are that, while specimens with higher ploidies are theoretically possible, there have been no records of these and, further, nQuire is only able to accurately detect ploidies up to 4x. If a specimen with higher ploidy existed, this should still be detectable across the many loci used and would result in an inconclusive estimate, thus ultimately being excluded from the study. As such, this cut-off provides a conservative dataset. Once loci were identified, I created a reference sequence by concatenating the reference genes used for the GoFlag 451 bait design for the most closely related species to *C. distans*, giving preference to *Gaga arizonica* sequences when available, and otherwise using *Notholaena montieliae.* I then aligned raw reads to the reference sequence using bwa (Li 2013), and sorted resulting reads using SAMtools (Li et al. 2009). The resulting BAM files were then uploaded to nQuire and denoised. This programme uses sequence alignments to make SNP calls that are then used to calculate the proportion of bases observed relative to each other for each site, which are then plotted to a histogram for all SNPs. We expect diploids to exhibit a normal curve with a peak centred at a 1:1 distribution of SNPs (50%); for triploids, we expect two normal curves with peaks at 33% (1:2) and 66% (2:1); and for tetraploids we expect peaks at 33%, 50%, and 66%. To further corroborate ploidy estimates, I also included sequences of a *Cheilanthes ecuadorensis* specimen for which a chromosome count exists (Sosa et al. 2021).

#### 2.2.3 Range mapping

The data for specimens with spore counts were uploaded to Quantum GIS 3.10 (QGIS 2017), where a map that plotted specimens coded by spore count was prepared (decimal degrees, projection: Equal Earth, coordinate system: WGS84). For visualisation, world administrative boundaries were downloaded from GADM (Hijmans et al. 2010. [continuously updated]).

In order to estimate whether the range size occupied by the different reproductive modes of *C. distans* were significantly different, I first calculated the total area (convex hull) for sexual and apomictic specimens in Quantum GIS. I then calculated bootstraps on these values with lmboot (Heyman 2019); these analyses and all others below were run in R version 3.6.0 (R Core Team 2017) and visualised with ggplot2 (Wickham 2016), except where noted.

### 2.3 Effect of spore size on dispersal

Spore sizes were mapped using Quantum GIS 3.10 (QGIS 2017) and a map prepared to visualise the data using the same parameters as in 2.2.3. In order to test whether smaller spores are able to attain larger ranges, I tested the correlation between range size and spore size. To determine spore range, I first split mean spore diameter data using three different break schemes: five quantiles or bins (each split equalling 20%), ten bins (each split equalling 10%), and twenty bins (each split equalling 5%). I then used Quantum GIS to calculate the total area (convex hull) for each bin in each of these schemes. The area values were then assigned to mean spore diameter values and linear regressions testing range to spore size were modelled. While the 20% split data found a significant relationship, all signal was lost in the other two splits. For this reason, I then visualised the raw data. Visual inspection revealed that the data were likely to be better explained by a parabolic regression and by smaller groupings (more bins). For this reason, I then created two additional splitting schemes of 15 bins (6.66% splits) and 25 bins (4% splits), calculated range area for these, and modelled parabolic regressions on the 10, 15, 20, and 25 bin splits.

### 2.4 Cataloguing spore forms and their effect on spore size and germination

While carrying out dissections to characterise reproductive mode and spore size, I noticed that there were many sporangia with irregular spores and that these irregularities fell into recurring patterns. This variation has not been characterised before, and may affect both dispersal (through spore size) and germination. For these reasons, I used the images captured in 2.2 to qualitatively sort sporangia into spore forms (Table 1; Fig. 1), with determinations for spore form made at the sporangium level. As such, the spore form determinations also transfer to the spores within that sporangium, given that they all resulted from the same meiotic event. Non-viable spores were those that had collapsed exospores, were overly transparent, or were not uniformly spherical (Hornych and Ekrt 2017). Sporangia with spores of similar size were judged to be those where all spores were within 15% in size to each other. The smaller spores in form E were often 70% or less the size of the other spores in the sporangium. Visual assessment of viable spores is most often clear cut, although there are occasional single spores that—although irregular in some way—seem like they may be viable (about 2%). In these cases, and since I made determination at the level of the sporangium, I was able to disregard individual spores and score spore form based on the rest of the spores. I summarised the results of this survey with a bar chart. I also calculated spore abortion index for apomictic plants by counting the number of aborted spores in the first image of 80 random specimens (to reach the recommended value of 1000 spores; Hornych and Ekrt 2017).

**Figure 1:**
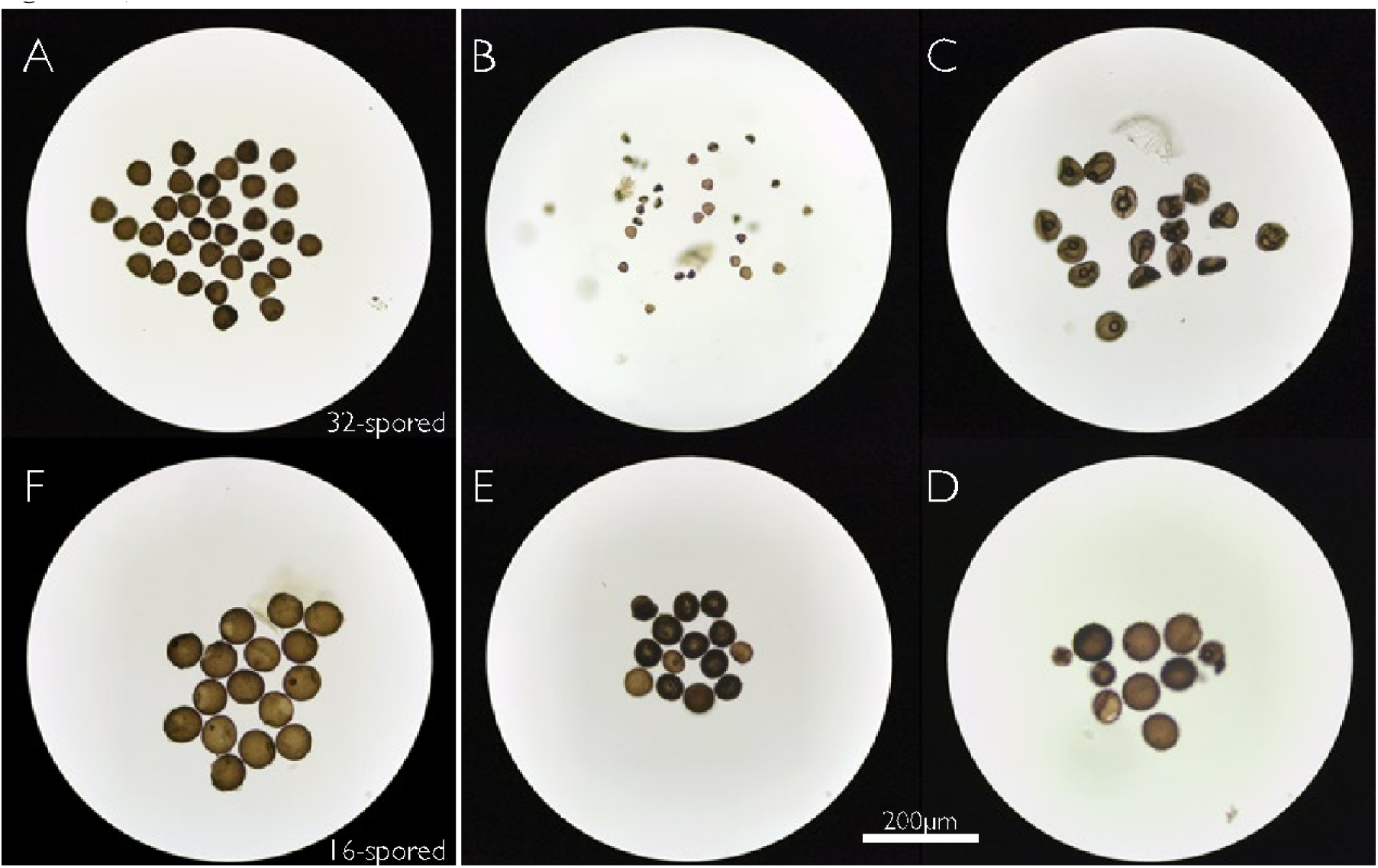
Spore variation in *Cheilanthes distans*. The left section shows well-formed sexual spores (32-spored sporangia; A; top) in contrast with well-formed apomictic spores (16-spored sporangia; F; bottom). The remaining images are labelled according to the spore forms outlined in Table 1.

**Table 1:** Spore forms and their hypothesised biological rationale.

| Code | Description | Biological Rationale |
| --- | --- | --- |
| A | 32 viable spores of similar size | Sexual diploid plant generating viable sexual spores |
| B | Non-viable spores of many sizes | Recent polyploid plant where traditional meiosis has broken down |
| C | 16 non-viable spores of similar size | Polyploid plant where non-division in meiosis I is established, but chromosomes are not sorted properly |
| D | Many sizes of spores but some seem viable | Apomictic polyploid where some chromosomes sort properly, but non-division in meiosis I is not established |
| E | 16 viable spores but 2-4 are smaller | Apomictic polyploid where both non-division in meiosis I and chromosome sorting are mostly established, with minor mistakes |
| F | 16 viable spores of similar size | Apomictic polyploid where both non-division in meiosis I and chromosome sorting are firmly established |

In a recent paper by Grusz and colleagues (2021), the authors posit that the shift from sexual reproduction to obligate apomixis must occur over a series of steps that need to become ‘locked in’. If the spore forms we observe (Table 1; Fig. 1) are steps along this pathway from sexual to asexual reproduction then, given that a single plant often shows more than one form, we might expect forms that are ‘near’ each other along the pathway (i.e. consecutive steps) to occur more frequently together (i.e. for forms within a plant to not be randomly distributed). Given that I have the data to assess this hypothesis, and that it may allow us to possible further understand spore formation dynamics, I scored each sample for presence or absence (0/1) for each spore form and tested for pairwise correlations between forms using corrr (Jackson et al. 2016) and using corrplot (Wei et al. 2017) to calculate confidence intervals.

Although some studies have looked at spore contents (Gemmrich 1977; Gómez-Noguez et al. 2016), and although spore forms have been briefly noted before (Manton 1950; Mehra and Singh 1957; Braithwaite 1964; Hickok and Klekowski Jr. 1973; Walker 1985), little is known about the effect of spore size and form on germination and viability. I aimed to capture a first glimpse into the relationship between germination, spore size, and spore form. For this experiment, only samples collected in the field in 2019 were used, to control for specimen age and treatment. Ten intact sporangia were removed from each of the six specimens available and were dissected, photographed, and annotated as outlined in 2.2.1. Once again, only viable spores were measured. In addition, I recorded both the total number of possibly viable spores (i.e. excluding spores with collapsed exospores, those overly transparent, or those anomalously shaped, but in this case including those with stable exospores and natural colour in unusual shapes) as well as the likely number of viable spores (i.e. this count excluded highly irregular spores and spores ruptured during dissection), along with the spore form for each sporangium. The contents of each individual sporangium were then transferred to an individual petri plate with Hevly’s by using a jet of water to wash the spores from the glass slide onto the plate, since individual spore transfer would likely rupture spores. Growth conditions were the same as those in 2.2.2. Growth of gametophytes and subsequent sporophyte development were monitored over 150 days, until all plants either reached the sporophyte stage or died from constrained conditions in the plates. Summary statistics were calculated for proportion of spores that germinated into gametophytes and proportion of spores that developed into sporophytes. Values were proportional because they were calculated at the level of the sporangium rather than for individual spores; I used the number of possibly viable spores for this calculation. The correlation between germination and spore size was tested using a logistic regression. Summary statistics for germination by spore form were calculated and visualised with boxplots. An ANOVA and subsequent Tukey test were run to test for significant differences in germination between forms.

## 3. Results

### 3.1 Locating the sexual diploids and characterising spore variation

#### 3.1.1 Spore data collection

A total of 416 herbarium specimens were examined. From these I obtained spore counts from 301 specimens (resulting in 740 spore per sporangium counts, averaging 2.5 sporangia per specimen) and spore diameter measurements for 197 specimens (resulting in 4825 individual spore measurements). A histogram of the total population distribution is shown in SI – Figure 1, and a histogram of the within plant variance is shown in SI – Figure 2. The remaining 115 specimens were either too young or all sporangia had already ruptured. Of the specimens with spore counts, seven had 32-spored sporangia (Fig. 1A)—making them presumed sexual diploids—and 294 had roughly 16-spored sporangia (Fig. 1C–F). I also catalogued spore form for 299 specimens, the forms of which are depicted in Figure 1; these results are discussed further below in 3.3.

**Figure 2:**
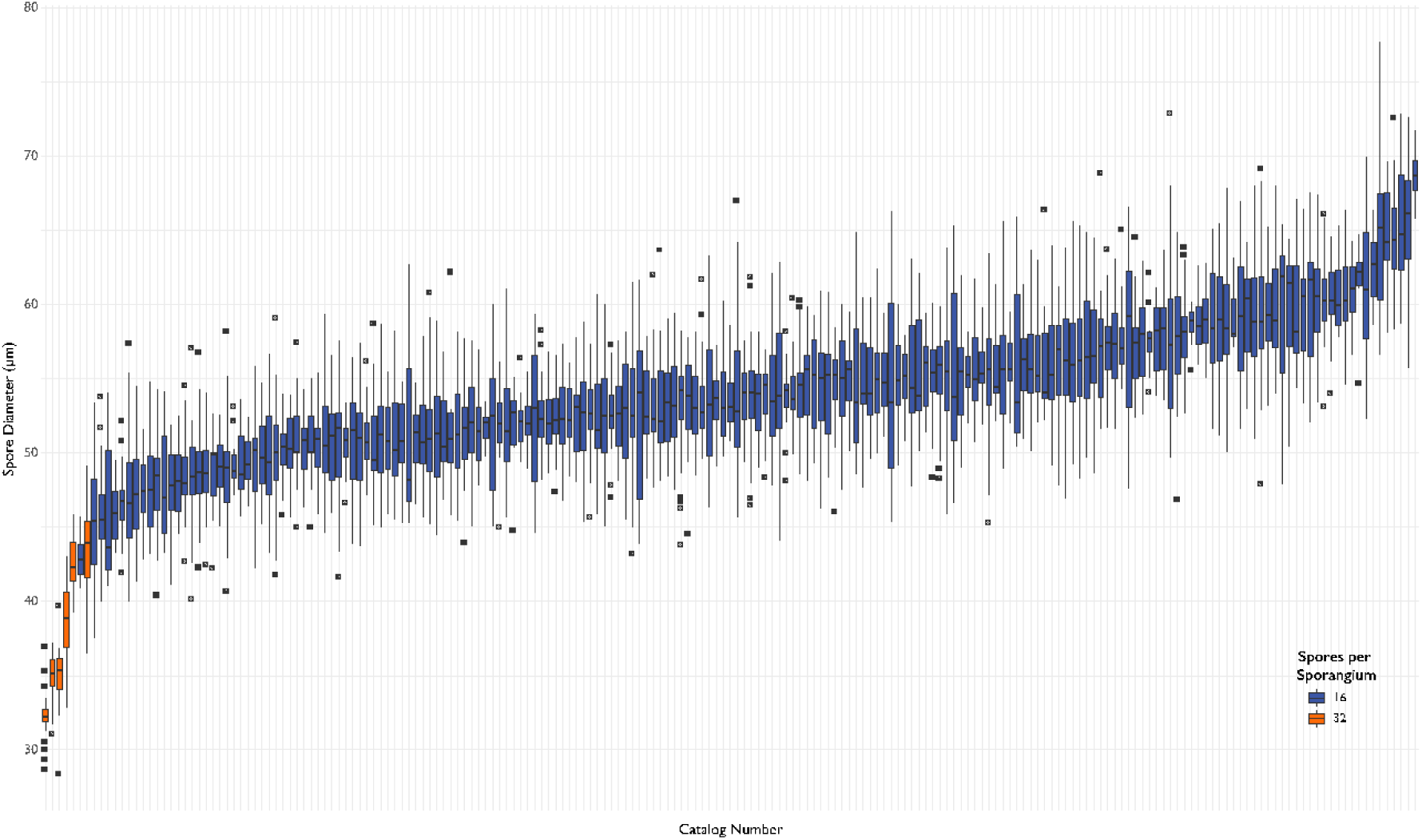
Boxplot of spore diameters for *Cheilanthes distans*. Orange boxplots on the far left are sexual specimens, and blue boxplots are apomicts.

#### 3.1.2 Ploidy estimation

Gametophytes for apomictic specimens developed after 11–25 days; they developed sporophytes apogamously after 35–55 days. I was successfully able to grow mature sporophytes from nine apomictic specimens. Sporangia began to develop after about two months. Although they produced many sporangia, due to the complications of the COVID-19 pandemic only a single chromosome squash was obtained (NSW 851335), which showed the specimen to be tetraploid (the image is not reproduced as the chromosome squash was done by Michael D. Windham but is available upon request.) None of the presumed sexual spores germinated into gametophytes despite multiple attempts under a range of light and temperature conditions, most likely due to the age of the specimens. As such we cannot say with complete certainty that these specimens do reproduce sexually, but will hereafter be referred to as the sexual specimens as other proxies (in particular spore number per sporangium and ploidy) do align with this categorisation.

Spore diameter has been used as a proxy for ploidy by many workers (e.g. Sigel et al. 2011; Huang et al. 2006). However, the variation in spore sizes in *C. distans* appear to be nearly continuous (Fig. 2), in contrast to the clear breaks observed in other species (e.g. Beck et al. 2010; Sigel et al. 2011). As such, it is not possible to visually determine which spore sizes correspond to which ploidy levels. Nevertheless, it is apparent that all sexual specimens do have smaller spores—as is expected by theory (Barrington et al. 1986).

I was able to obtain ploidy estimates for 18 of the 23 *C. distans* specimens for which I have sequence data; the remaining samples had either too little signal or equivocal results. Both calibration points (i.e. the specimens for which chromosome squashes exist) were accurately estimated by nQuire. I plotted spore size distributions for the specimens for which I was able to estimate ploidy (Fig. 3). The diploids correspond to the presumed sexual specimens and the above trend is corroborated, where these once again have the smallest spore sizes. While the tetraploids tend to have larger spore sizes, I find that one tetraploid specimen has spore sizes well within the triploid range.

**Figure 3:**
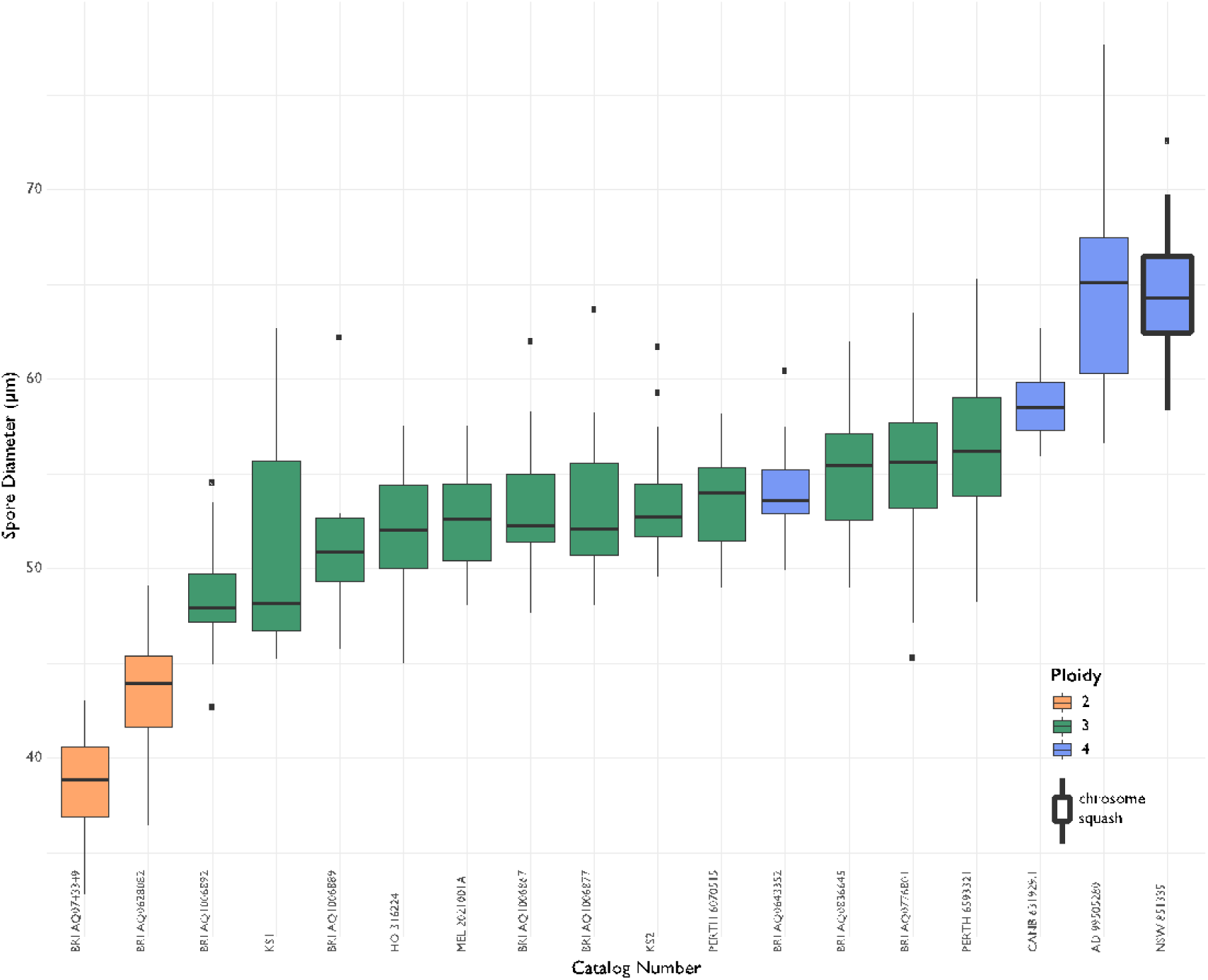
Boxplots of spore diameters for *Cheilanthes distans* specimens with estimated ploidies. Orange boxplots are diploids, green boxplots are triploids, and blue boxplots are tetraploids. The specimen on the far right (thick outline) has a chromosome squash that confirms it as a tetraploid.

To further test the visual trends observed in Figure 3, I ran an ANOVA on the grouped spore diameters for each ploidy level. Since the results were significant (p < 2e^-16^), I then ran a Tukey multiple comparisons test, which found all groups to be significantly different from each other (Fig. 4). However, given the results observed in Figure 3, it is nevertheless not possible to establish clear-cut ranges in spore size that correspond to ploidy levels. Instead, it seems like there are distinct yet overlapping spore size distributions.

**Figure 4:**
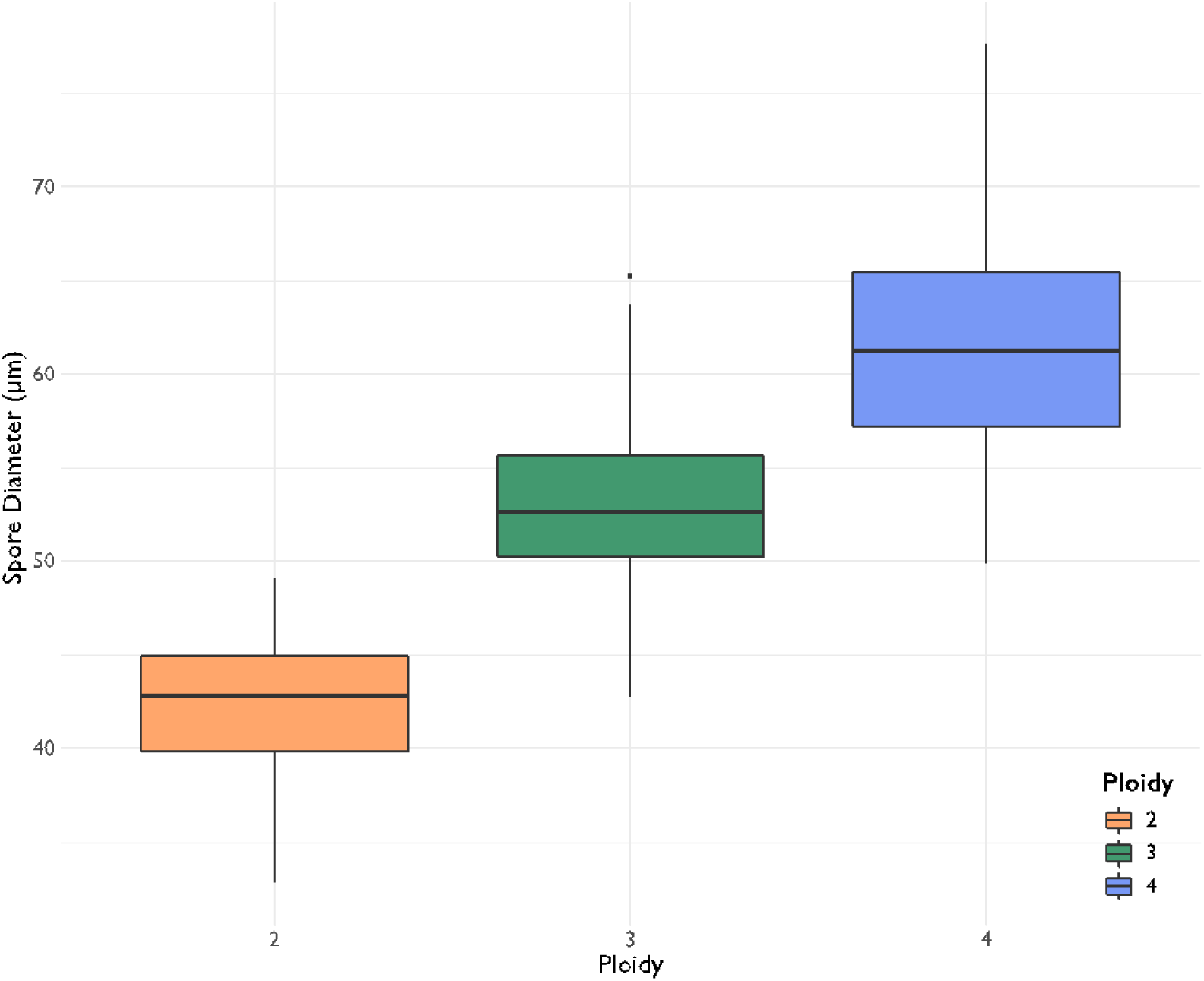
Boxplots for the ANOVA analysis between ploidy and spore diameter. All groups were found to be significantly different from one another.

#### 3.1.3 Range mapping

The range map for *C. distans* shows a clear example of Baker’s rule (Baker 1967), where the range of the apomict (which are inherently self-compatible) is far larger than that of the sexual specimens (which may not be self-compatible) (Fig. 5). The sexual specimens are mostly located in eastern Australia, with a single outlier located on the western coast. This outlier specimen is also unusual because none of the spores on the plant appeared viable, although perhaps they were simply immature. There are no other specimens collected in that region.

**Figure 5:**
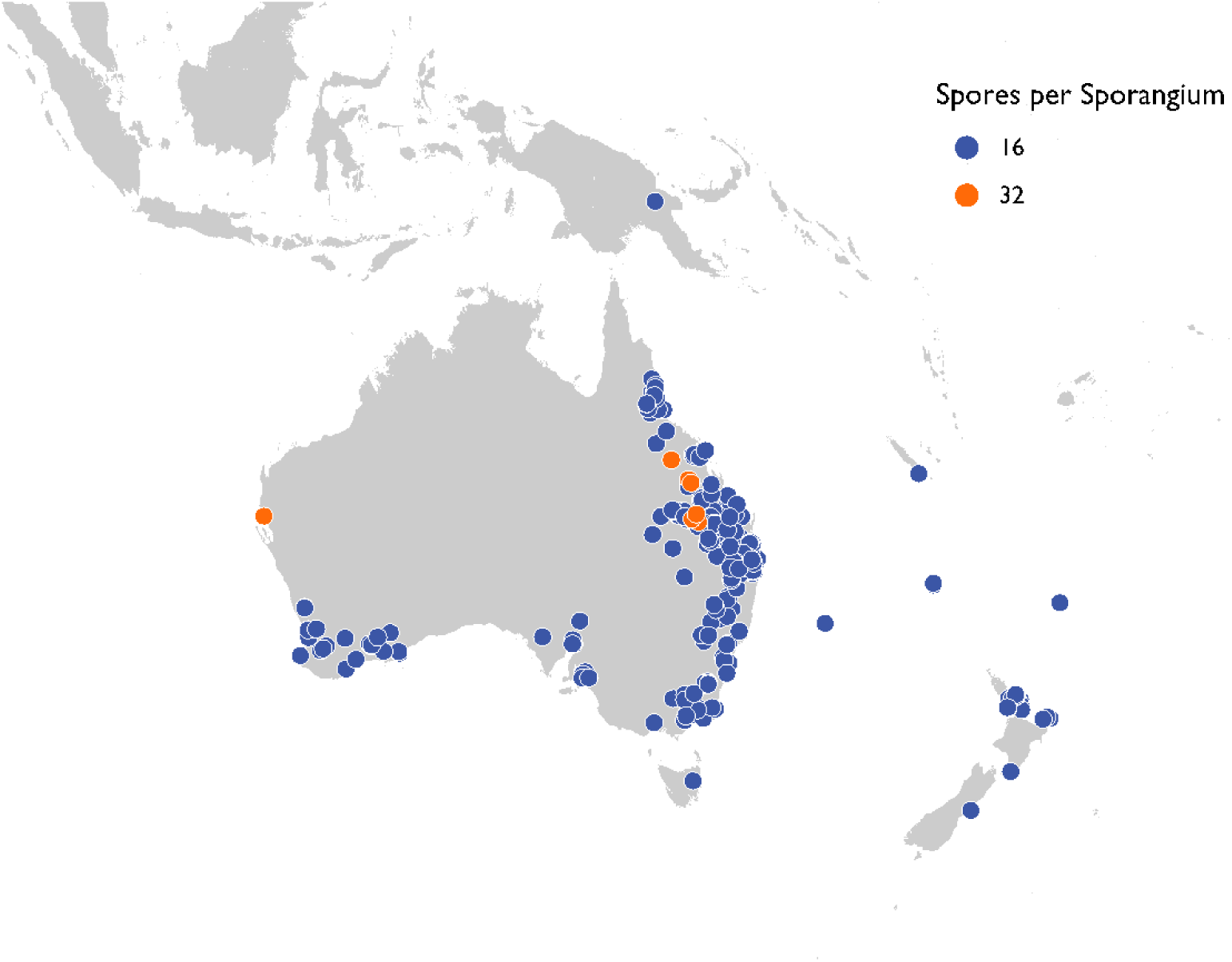
Range map for *Cheilanthes distans* spore counts. Blue dots represent apomictic specimens (16 spores per sporangium) and orange dots represent sexual specimens (32 spores per sporangium).

Because specimen sampling was done without knowledge of where sexual specimens might be found—if at all—and considering the relatively extensive sampling of this study, the sampling can be considered fairly random and the range for sexual specimens is thus likely to be similar to that found here. Further, the bootstrap analysis comparing the range sizes between sexual and apomictic specimens, and which takes into account differences in sample size between the two groups, found that their ranges are in fact significantly different (p=0).

### 3.2 Effect of spore size on dispersal

Geographic visualisation of spore size data show some structure in spore size, although it is not possible to outline clear trends from this map (Fig. 6). One can notice, however, that the smallest spores appear limited to a north-eastern Australia range, that the largest spores appear limited to a southern Australian range, and that medium spores seem most widely distributed.

**Figure 6:**
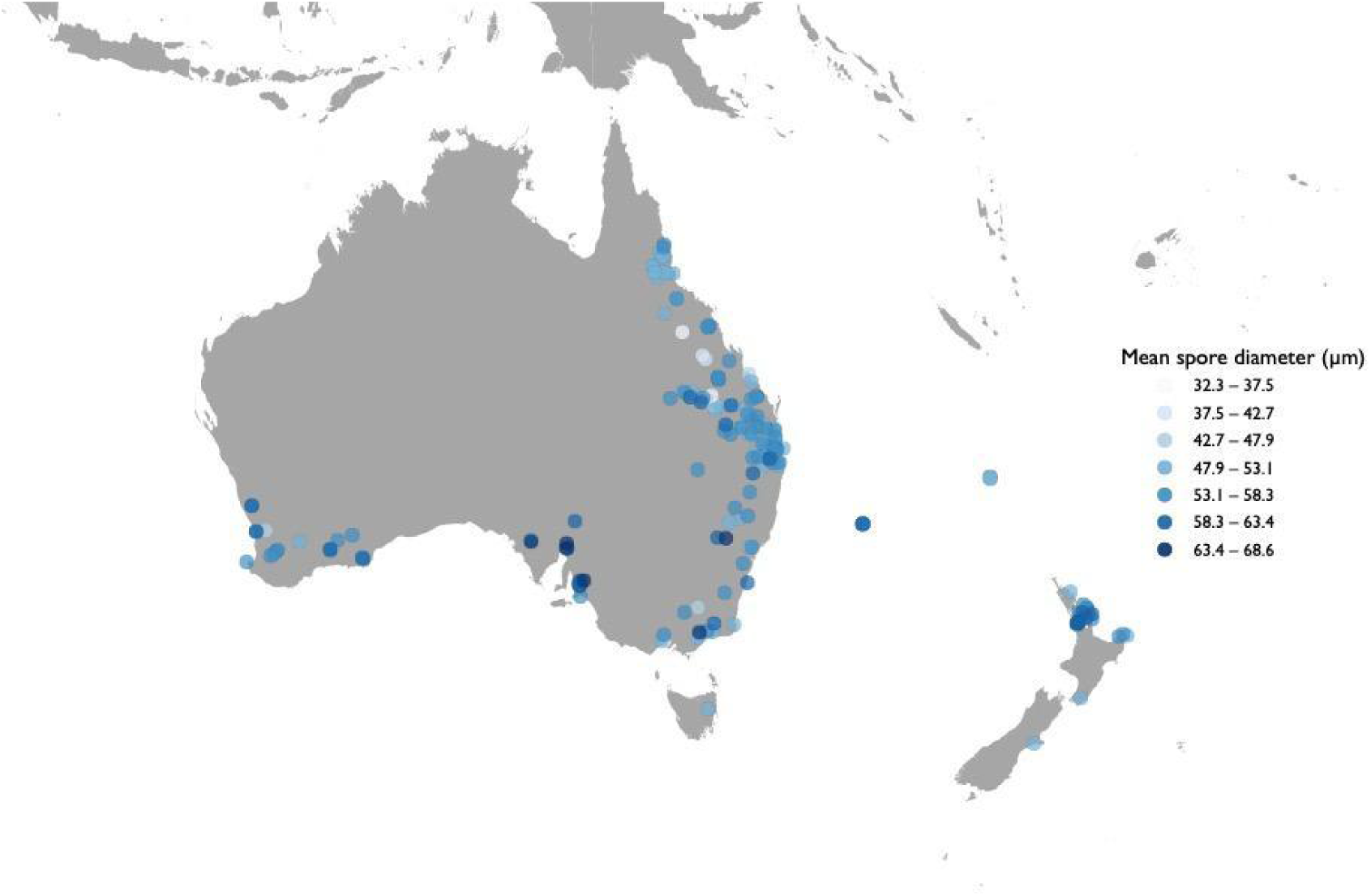
Map showing the distribution of *Cheilanthes distans* spore sizes. Lighter blue shades correspond to smaller spores and darken gradually into navy as size increases.

To explore these trends statistically, I then carried out regression analyses. The attempts to identify correlations between spore size and range size resulted in unexpected patterns, which were explored in two phases. Data for the splitting schemes used (20%, 10%, 6.66%, 5% and 4%) can be seen in Figure 7. Of the initial linear regressions for the three original splitting schemes (20%, 10%, and 5%), only the 20% splits data found a significant relationship between range area and spore size of functional spores, with an adjusted R^2^ of 0.353 (p<2e^-16^; Fig. 8A). Surprisingly, however, this relationship was completely lost for the other two linear regressions (Fig. 8A). Upon visual inspection of the raw values, I realised that the data were closer to a parabolic distribution and that the variation was best captured with the smallest of the three splits (Fig. 7). For this reason, I then did follow-up parabolic regressions using two additional splitting schemes, ultimately testing parabolic regressions for 10%, 6.66%, 5%, and 4% splits. Most parabolic regressions were highly significant (p≤1.84e^-05^; Fig. 8B), the sole exception being the 10% splits data (p=0.6). Of these, the best model was the one that used 4% splits since it had the highest adjusted R^2^ (equal to 0.123) and the lowest residual error (equal to 124.5). Nevertheless, this model predicted range size values for the smallest spores that are considerably negative. For this reason, and considering that all parabolic relationships show similar results, I have picked the 5% splits data to represent the ‘best’ model as a compromise between statistical results and realistic biological scenarios; for this model, the R^2^ equals 0.083 and the residual error equals 148.5 (Fig. 9).

**Figure 7:**
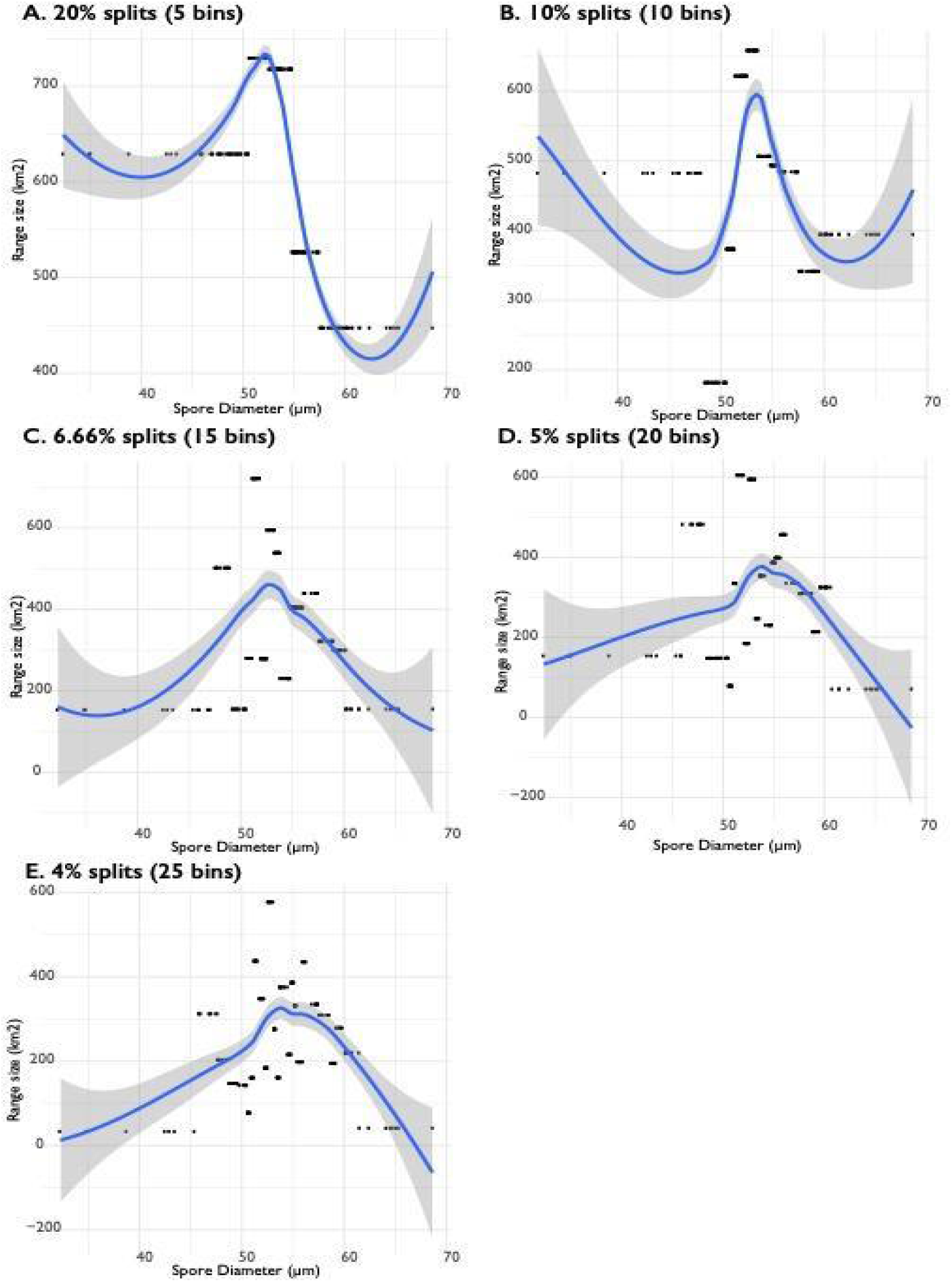
Data for spore diameter to range area, plotted by bins (bins are visible as dot clusters on the y-axis), with smoothed blue lines that track the data to aid visualisation. A) 20% splits B) 10% splits C) 6.66% splits D) 5% splits E) 4% splits

**Figure 8:**
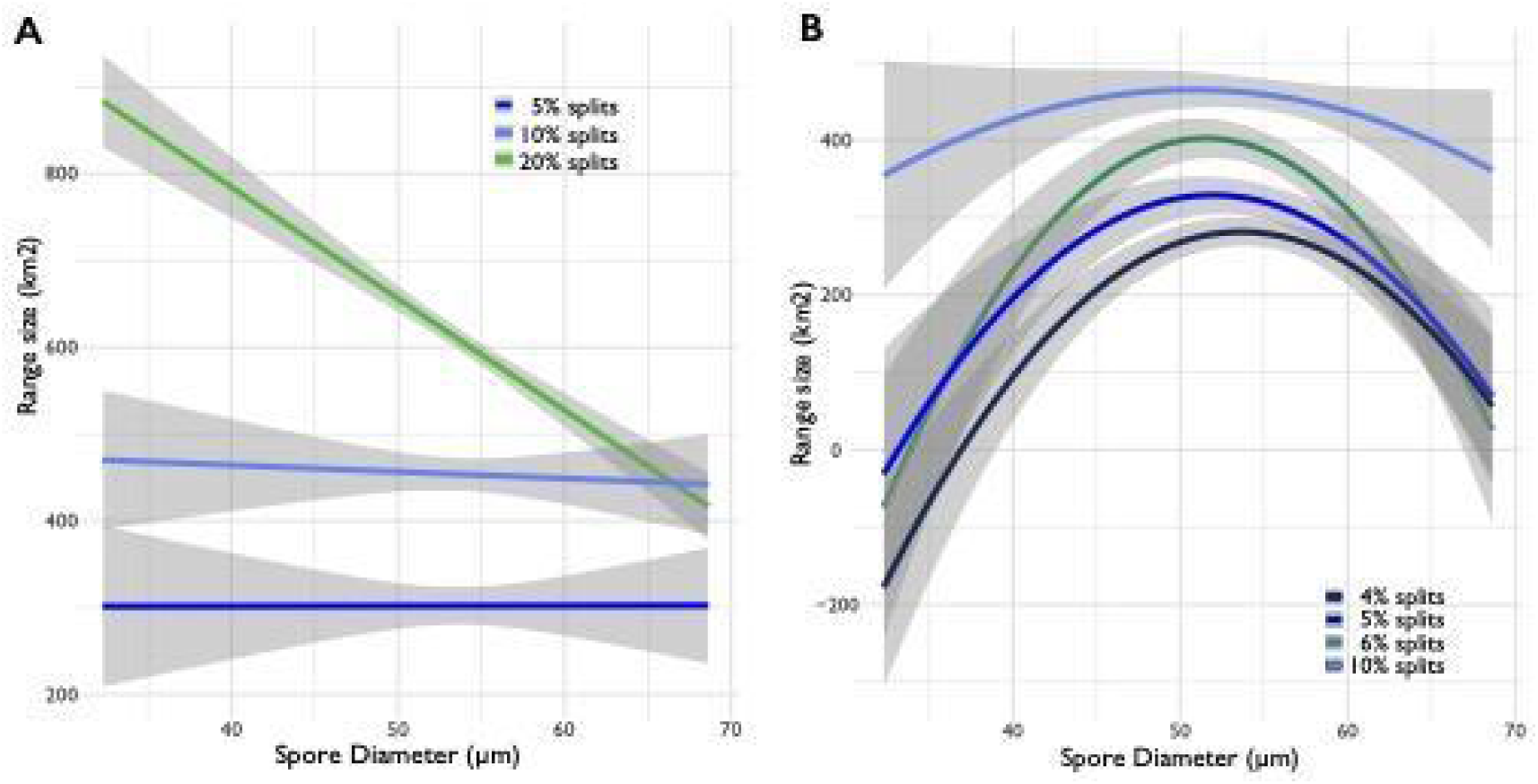
Regression tests for range area to spore size correlations. Colours are kept consistent across panels and with Figure 9. A) Linear regressions B) Parabolic regressions

**Figure 9:**
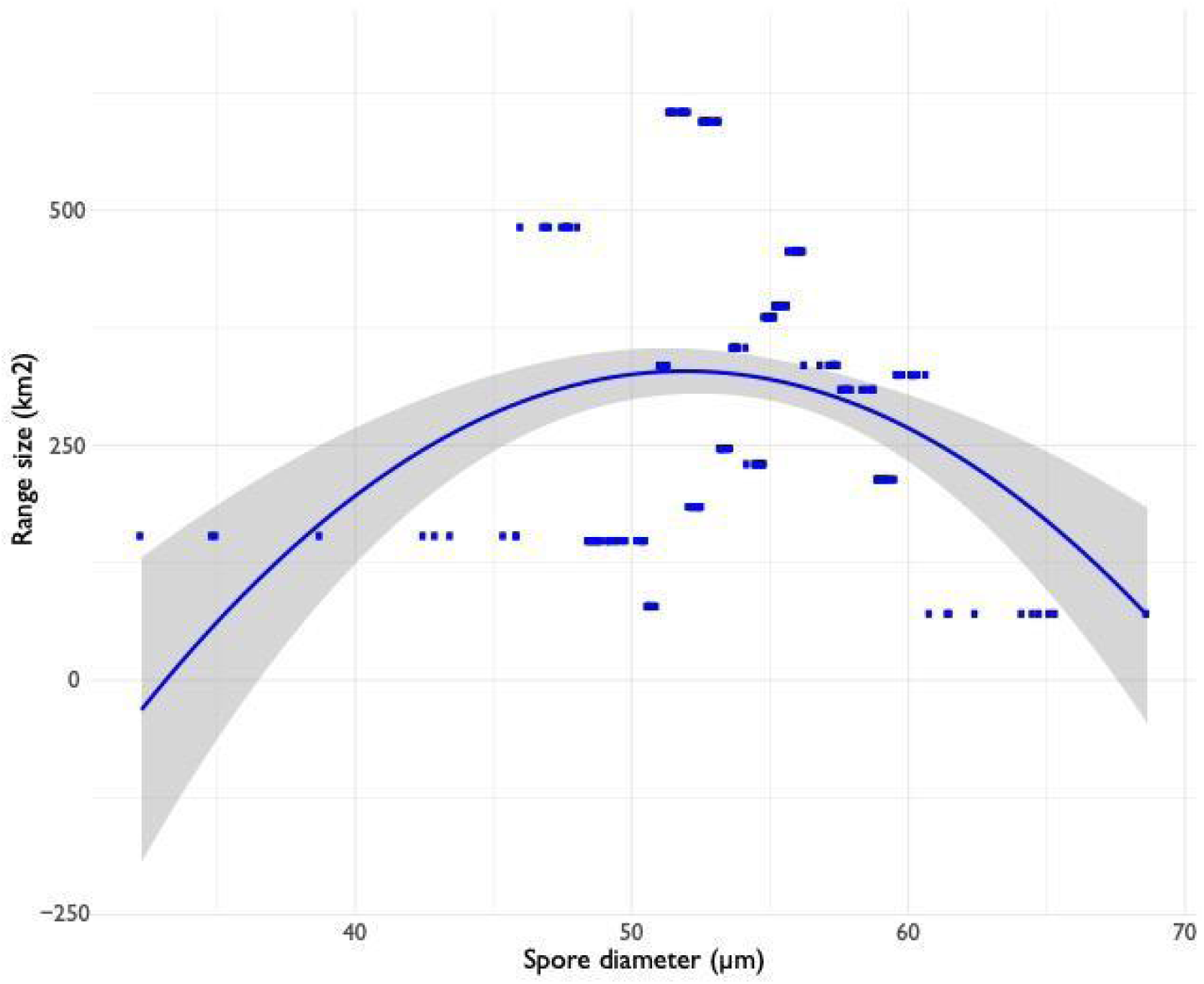
Best fitting regression for the relationship between range area and spore size, which uses 5% splits. The regression coefficient on *x* is 91.336 (S.E.=20.67; p=1.67e^-5^) and that on *x^2^* is –0.871 (S.E.=0.198; p=1.84e^-5^).

### 3.3 Cataloguing spore form and their effect on spore size and germination

A total of 299 specimens were catalogued for spore form, resulting in 737 total characterisations. The spore forms are characterised visually in Figure 1. The most frequent form were sporangia with 16 viable spores of similar size (form F; Fig. 10), followed by 16-spored sporangia with 2-4 smaller spores (E). Sporangia with 32 viable spores (A) and those with non-viable spores of many sizes (B) were the least frequent. The overall percentage of aborted spores out of 1000 was 23.8% (otherwise known as spore abortion index as per Hornych and Ekrt (2017)).

**Figure 10:**
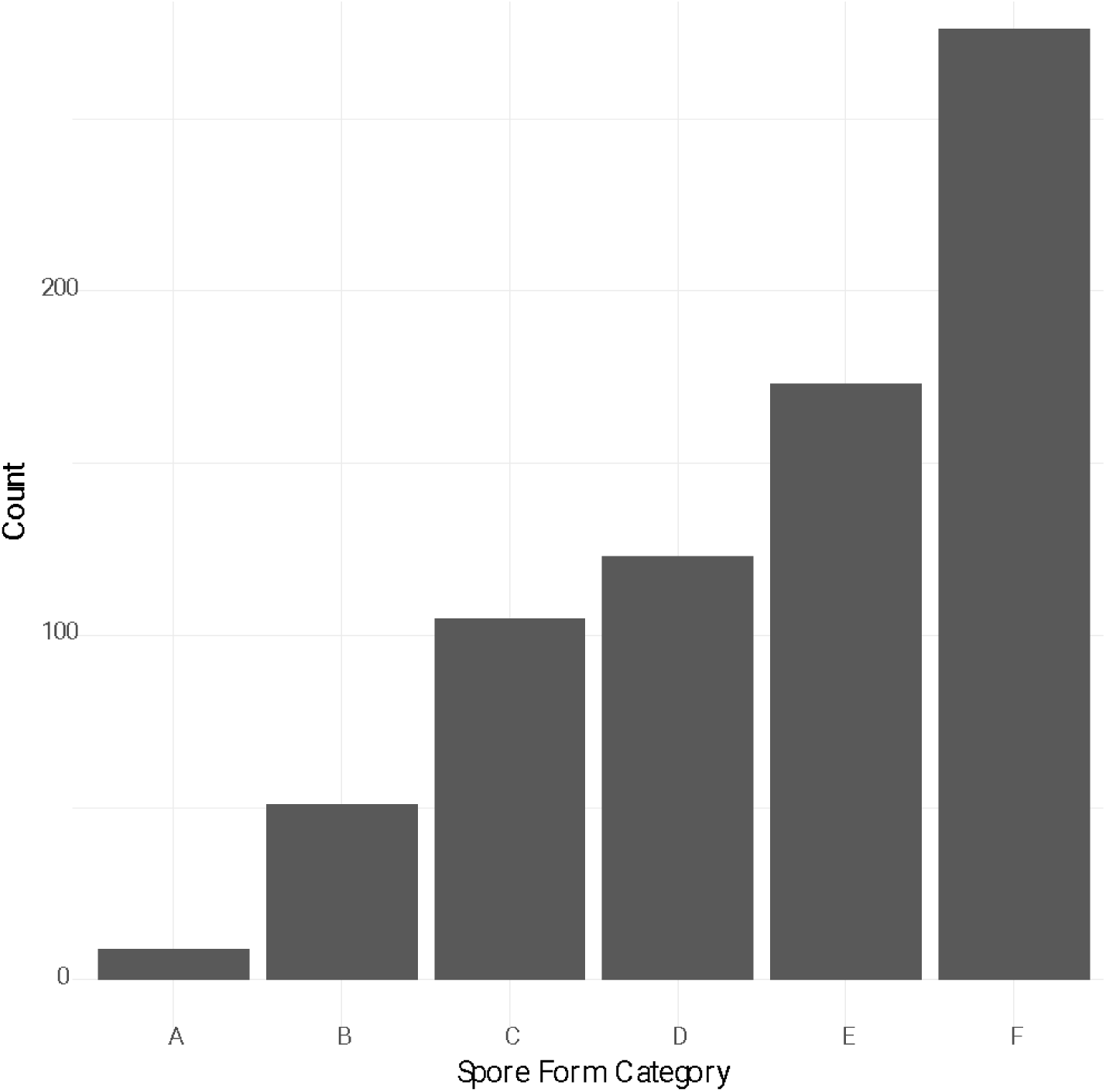
Bar plot of net counts for each of the spore forms in *C. distans*.

The correlation analysis between spore forms within a plant found that most forms are negatively correlated to each other (SI – Table 1). These values are quite small: the strongest negative correlation is between forms B and F (–0.34), while the strongest positive correlation is between forms A and B (0.09). Further, the confidence intervals are quite wide, most of them around ± 0.8, and none of the correlations are significant. In order for us to conclude that the spore forms are steps along a pathway, we would need to see non-recurring positive correlations between spore forms (such that we could deduce a sequence). Given that we almost exclusively see negative correlations and that they have such low confidence, it does not appear that the spore forms I catalogued are in fact steps along a pathway. Given the null results resulting from this analysis, they are not explored further.

I was able to obtain 10 samples per specimen for the germination trials with the exception of one specimen where I was only able to find eight intact sporangia; the sporangia ranged from having 0–16 viable spores, with a median viable spore count of 11. All the specimens sampled were apomictic. Of 559 spores scored visually as possibly viable, 253 developed into gametophytes and 230 of them grew to sporophytes.

The logistic regression for proportional growth to gametophyte relative to mean spore size was statistically significant, with a regression coefficient of 0.13 (p=0.05) and a standard error of 0.07. Values for the regression testing the proportion of spores that grew to sporophyte resulted in similar values: a significant relationship was found with mean spore size (regression coefficient of 0.12, p=0.05, standard error 0.07). Both regression lines are plotted in Figure 11.

**Figure 11:**
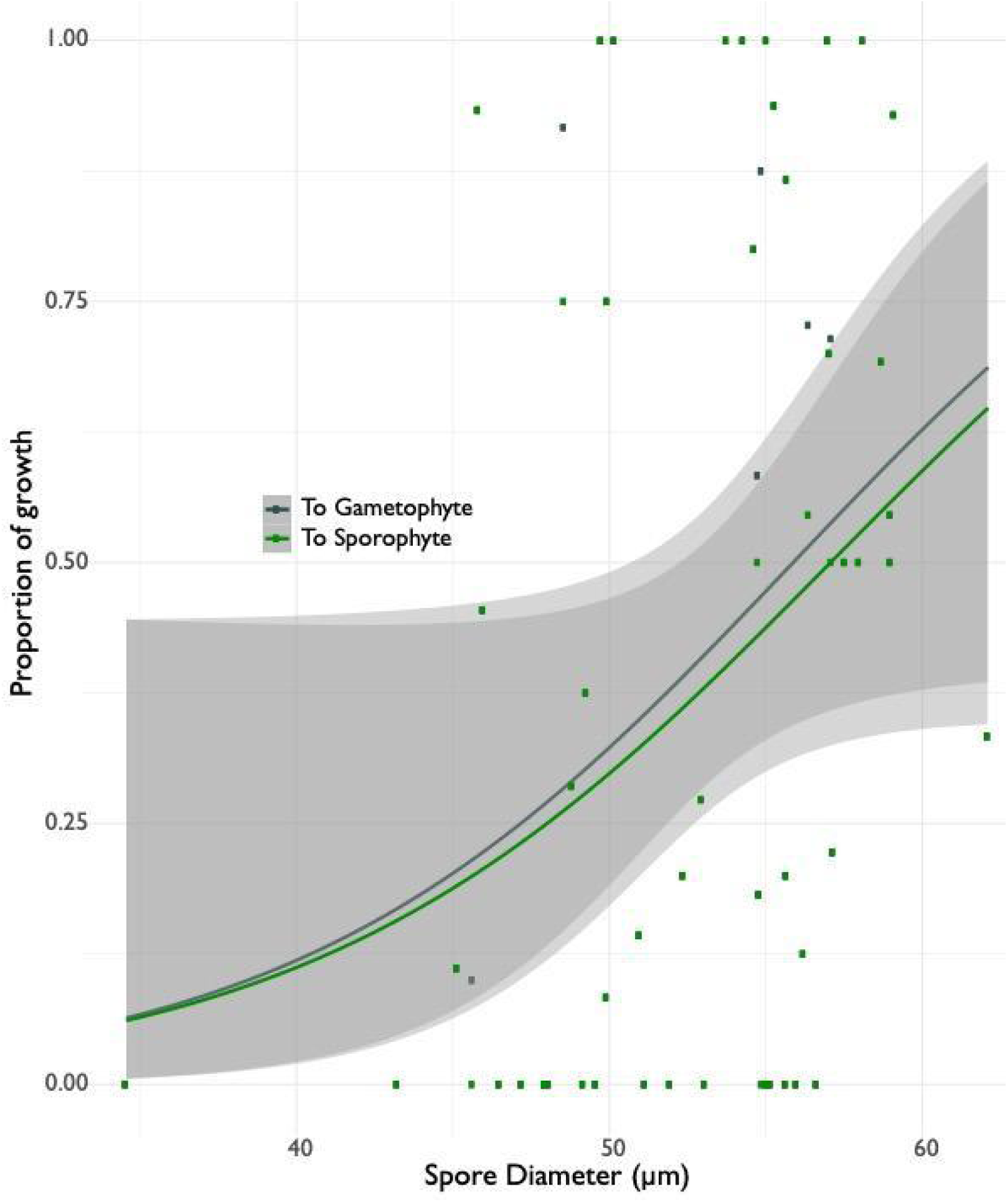
Logistic regression between mean spore diameter and proportional growth to either gametophyte (dark grey) or sporophyte (green).

As with other metrics, I looked only at viable spores and their development to sporophyte and found growth increase across spore forms (Fig. 12; SI – Table 2 lists raw numbers and growth to gametophyte data, which show a similar pattern). The forms have been arranged to follow the order outlined in Table 1, but it is clear that proportions increase as we move right along the forms. This is likely due to the fact that there are on average more viable spores available in these categories. The highest mean growth to sporophytes was seen in F form sporangia (i.e. sporangia with 16 well-formed spores) followed by forms D and E. It is likely that some of the loss of germination seen—particularly for form F where all spores should be functional—could be due to damage of spores during dissection, as spores are often ruptured even despite careful handling and this damage is not always easy to detect under light microscope examination. It is also worth noting that because the specimens available for the germination experiment were all apomictic, there were no spores of form A available to include in this part of the study. While the ANOVA found significant differences between the spore form groups (p=0.013), the sample size was too small for the Tukey test to be able to detect significance when doing pairwise comparisons.

**Figure 12:**
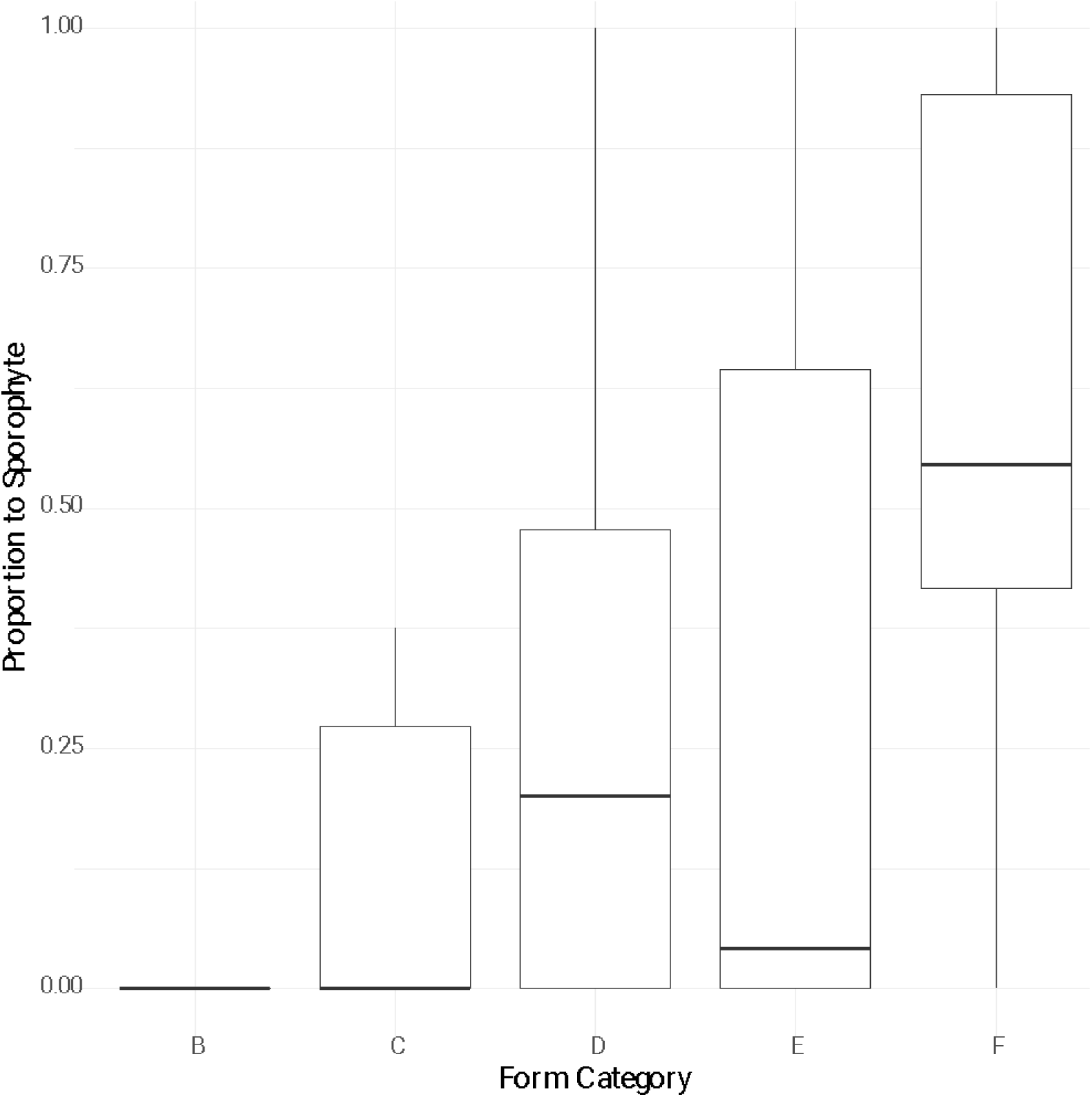
Boxplot for proportional growth to sporophyte by spore form. Note that no A form sporangia were found amongst these specimens, so this form could not be tested.

## 4. Discussion

### 4.1 Locating the sexual diploids and characterising spore variation

Understanding the biological processes that drive—or limit—dispersal and range size in organisms remains of perpetual interest in evolutionary biology. Numerous studies have found that self-compatible and asexual organisms can establish populations over greater distances than their outcrossing, sexual counterparts (Coughlan et al. 2017; e.g. Hörandl 2006; Hörandl et al. 2008; Marchant et al. 2016; Wickell et al. 2017). Past studies have also identified this trend in xeric-adapted ferns (Beck et al. 2010; Sigel et al. 2011; Dyer et al. 2012), but the geographic ranges of their studies were smaller than the present one, and none were located in the southern hemisphere or in Australasia. I was interested in testing whether larger ranges for asexual organisms would hold true for a xeric-adapted fern with a patchy habitat and spread over a larger geographic range, as it was unclear whether the environmental variation might be too great at the above-continental scale for this trend to hold, or whether xeric environments might—at this scale and in this region—be harsh enough that sexual reproduction could be favoured as a reproductive strategy, as studies in other groups have found that diploids can fare better in some harsher conditions (García-Fernández et al. 2012; Tanaka et al. 2014).

The survey of *Cheilanthes distans* was able to locate the presumed sexual progenitors—a lineage new to Western science—to the apomictic lineages of this species. Despite extensive sampling, only six viable sexual specimens were located, found over a relatively narrow range (Fig. 5). This pattern falls in line with past studies and ultimately supports Baker’s rule, which posits that self-compatible and asexual organisms will be able to spread over larger distances, as they don’t need mates to establish new populations (Baker 1967). In the case of ferns, this rule might theoretically be slightly relaxed, since single spores developing into sexual gametophytes may be self-compatible through gametophytic selfing and thus be able to produce offspring with only a single propagule, barring any lethal mutations in a particular gametophyte (Haufler et al. 2016; Sessa et al. 2016; Klekowski Jr 1982). Yet, even given the long-term advantages allowed by sexual reproduction (Stebbins 1957; Muller 1964; Darlington 1946), asexual *C. distans* are able to attain a larger range, possibly also pointing to difficulties in self-compatibility in this species when resulting offspring are completely homozygous. It is worth noting that sexual and apomictic lineages overlap in their range, so that this species does not exhibit geographic parthenogenesis per se (since geographic parthenogenesis occurs only when sexual and apomictic ranges are distinct). It remains worth exploring how flexible apomicts can be in relation to environmental conditions and whether we can detect environmental variables that constrain sexual lineages to their present range. Apomictic triploids are apparently much more common in *C. distans* as compared to tetraploids (Fig. 3). The reasons why triploids manage to be more successful than tetraploids are also of considerable interest, since they both should be able to reap the benefits of asexual reproduction. It is possible the lower numbers of tetraploids may at least in part be linked to spore size, which is discussed further in 4.2 below.

It was unexpected to find spore size distributions that were as continuous as those observed (Fig. 2), given past observations that reported discrete spore size groupings. However, a sampling of spore diameters has not been done with such a large number of samples before. The size variation found is similar in distribution to that observed in pollen size (Kelly et al. 2002; Jürgens et al. 2012), so the spore size variation found is perhaps ultimately not surprising, although we do observe slightly higher total population variance in the spore data (37.08 compared to 12.96–27.04 in pollen in Kelly et al. (2002) and Jürgens et al. (2012); SI – Fig. 1), in addition to high within plant variance (SI – Fig. 2). This higher variance may in part be due to the pathway that produces spores in apomictic *C. distans:* first division restitution meiosis has been reported to be prone to unequal divisions and mistakes in the distribution of chromosomes during sporogenesis (Braithwaite 1964; Walker 1985; Mehra and Singh 1957; Hickok and Klekowski Jr. 1973). Moreover, all previous apomictic fern studies that have searched for spore size breaks as proxies for ploidy have studied groups with spores produced through premeiotic endomitosis, which is reported to be relatively stable and mostly result in viable spores (Grusz 2016; Grusz et al. 2021). In fact, I find that variation in spore size in *C. distans* is such that we cannot clearly delimit proxies for ploidy based on spore sizes (Fig. 3), even as there is a tendency for higher ploidies to have larger spores (Fig. 4). Indeed, this trend has also been found in recent work by Barrington and colleagues, who noted there is significant variation in spore size between fern species of the same ploidy (Barrington et al. 2020). Regardless, spore size variation may ultimately be fertile ground upon which selection can act through the dual processes of dispersal and establishment, as will be discussed below.

### 4.2 Effect of spore size on dispersal

Many studies across a range of systems have shown that smaller dispersal units—be they spores, pollen, or seeds—tend to travel farther (Norros et al. 2014; Muller-Landau et al. 2008; Wilkinson et al. 2012 and citations therein). In the case of ferns, the higher abundance of fern to spermatophyte species on isolated oceanic islands (compared to this same proportion in continental areas) is testimony to their dispersal ability (Tryon 1970; Smith 1972; Peck et al. 1990). However, past work has been limited in the scale at which it tested dispersal, or conversely has been solely theoretical. To my knowledge, the present study is the first to empirically test the effect of spore size on dispersal over such a large area (approximately 18 million km^2^). However, I find what is likely to be an interesting interplay of biological forces (Fig. 9) that is more complex than simply smaller things travelling farther and therefore having the largest ranges.

The consistently found and well-supported parabolic model suggests that there may be more than one main driver when considering the interaction between range size and spore size, since it is individuals with medium-sized spores that are found to attain the largest ranges. This relationship holds even when we remove sexual specimens (data not shown), so there must be drivers simply beyond reproductive mode that are causing this pattern. We need only look at the results from the germination experiment (3.3) to posit a hypothesis: spores of larger sizes were found to have higher germination, so it may be that the parabolic relationship between spore size and range results from two opposing processes: smaller spores disperse farther (as per theory and past studies: Gregory 1945; Norros et al. 2014; Hussein et al. 2013; Tesmer and Schnittler 2007) but larger spores have greater survival to sporophyte. This trade-off ultimately leads to individuals with spores of intermediate sizes having the greatest ranges. In particular, spore provisioning may be especially important given the harsh xeric environments where these ferns grow: data from seeds show that larger seeds are especially important for seeding survival when they germinate in harsher conditions (Bruun and Ten Brink 2008; Westoby et al. 1992; Jurado and Westoby 1992, which looked specifically at Australian desert plants). Although the spore volume of *C. distans* are small compared to most seed plants, most provisioning in ferns is done through lipids (Gemmrich 1977; Ballesteros and Walters 2007), which are lighter, have a longer shelf life, and are more energy dense than the carbohydrates usually provided by seeds (Gemmrich 1977; Gómez-Noguez et al. 2016), so that spores may indeed pack more provisions than we might expect.

Past studies have indeed also found that there are trade-offs between dispersal ability and life-history traits related to size such as seed/spore germination and young sporophyte growth and survival (Norros et al. 2014; Eriksson and Jakobsson 1999; Bruun and Ten Brink 2008; Chrtek et al. 2018), but this study is the first to do so in ferns and only the second to look at trade-offs involved in plant spores (Löbel and Rydin 2010). The fact that propagule size appears to be broadly consistently subject to trade-offs between dispersal and survival in all these studies—despite the fact that spores germinate and reproduce quite differently from seeds—hints at the fundamental nature of certain reproduction-related trade-offs.

Given our current understanding of first division restitution sporogenesis as relatively error-prone, it remains worth exploring the heritability of spore size, as well as testing through direct experimentation whether the apparent trade-off detected by our model is in fact at play. Given the large range over which this species exists, it is possible that the large variation in spore size observed—often within a single plant—might enhance fitness by producing spores that are ‘best suited’ to a wide range of environmental conditions—for example generating larger spores that might be the only ones to survive in harsh conditions—thus maintaining spore variation as a ‘trait’ (Stearns 1989). In fact, large numbers of offspring with considerable size variation has often been observed as a bet-hedging strategy adopted by species faced with unpredictable environments (Olofsson et al. 2009; Morrongiello et al. 2012; Simons 2007). Although the predictive power of the 5% splits parabolic model is somewhat low (adjusted R^2^=0.08), it is valuable as a first step in gaining a greater understanding of the trade-offs that influence fern spore sizes.

### 4.3 Cataloguing spore form and their effect on spore size and germination

Past workers have characterised sporogenesis as acting seemingly independently in each sporangium, such that each meiotic event can result in different forms and numbers of spores (Ekrt and Koutecký 2016; Mehra and Singh 1957; Bell 1960). As a result, sporogenesis can vary greatly within a plant. In fact, the extensive spore survey carried out in efforts to locate the sexual lineage of *C. distans* provided a catalogue of variation in spore forms that may arise from modifications to meiosis (Table 1). The variation in forms observed in *C. distans* would seem to support this hypothesis, especially since the correlation analysis did not reveal trends in the distribution of spore forms within a plant. Further, the within plant spore size variance can be quite high, with some samples even having a variance greater than 40 (SI – Fig. 2). My survey results show extensive variation in fern spores that may play a role in dispersal and survival, and that is beyond what has been described in the past, where spores were thought to be either clearly well-formed or otherwise abortive, and to show only narrow variation in spore size (e.g. of many: Manton 1950; Bell 1960; Hornych and Ekrt 2017; Sigel et al. 2011).

If we focus on apomixis as a reproductive strategy, in order for it to become obligate in a lineage several modifications need to both arise and become ‘locked in’. Grusz and colleagues have posited a hypothesis for the steps needed for a change from sexual reproduction to obligate apomixis to occur, beginning with facultative unreduced spore production that results in obligate unreduced spore production, which leads to apogamous reproduction that eventually and ultimately becomes fixed (Grusz et al. 2021). It is of interest whether we can locate the intermediate stages outlined in their paper, and whether the spore forms observed in *C. distans* may be these intermediate steps. If it were the case that certain modifications to meiosis need to occur before others—forming a ‘pathway to obligate meiosis’—we might expect steps along said pathway to occur more frequently together (i.e. in the same plant). My results, however, detected only two positive correlations among spore forms present within a plant out of fifteen comparisons. As such, these results are insufficient to posit a ‘pathway’ between all forms. Most of the correlations observed are negative ones between F and all other spore forms, followed by E and all other spore forms. This may suggest that once correct (F) to near-correct (E) sporogenesis is achieved in a plant, it will then occur most frequently, with other spore forms remaining possible although rarer. It is worth highlighting that our sample size per plant is quite small (three sporangia per plant). It is possible that larger sample sizes may be needed to detect these correlations, that correlations are happening at the level of the population rather than the plant, or simply that the hypothesis I proposed is incorrect.

The spore abortion index for *C. distans* at 23.8% is in the higher end of values reported for other apomictic ferns by Hornych and Ekrt, who find two groups: one averaging 13.74% and another averaging 24.8% abortion (Hornych and Ekrt 2017). While the data available do not allow us to test for statistically significant difference between these two abortion index groups, my findings nevertheless support past observations that apomictic sporogenesis via first division restitution seems on average more prone to mistakes (Braithwaite 1964; Walker 1985; Mehra and Singh 1957; Hickok and Klekowski Jr. 1973; Grusz 2016). Given apomictic spores propensity to mistakes, the fate of these spore forms with regards to germination and survival are of interest. Although several studies have noted the presence of spores that appear viable amongst aborted spores (Ekrt and Koutecký 2016; Hornych and Ekrt 2017; Hickok and Klekowski Jr. 1973), or that have tested germination in specimens with notable abortion rates (Gomes et al. 2006), none of these studies has catalogued spore variation and correlated it with rates of germination. Despite a limited sample size, I was able to track spore germination over a range of spore forms and spore sizes (albeit with these data grouped at the level at the sporangium). I found that purportedly unviable spores—regardless of the spore form they arise from—can germinate and develop into sporophytes. Unsurprisingly, spore forms with more classically-viable spores (i.e. E and F) tended to have higher germination rates, but this may simply reflect that on average there are higher numbers of viable spores in these sporangia (since germination values were calculated using only viable spore counts). As such it doesn’t appear that certain spore forms are intrinsically viable or unviable, but rather that viability is determined by the fate of each spore resulting from any given meiotic event. These data point to the important role that rare or relatively rare events can have in shaping evolution, as a mostly abortive sporogenesis need only generate a single viable spore for a subsequent generation to become possible. In fact and most surprisingly, we find that spores that might be categorised visually as unviable (in this case, due to their non-spherical, faceted shape) are in fact able to germinate and develop into sporophytes (see Fig. 12—spore form D). Although visual estimation of viable spores remains valuable and broadly accurate, these results point to its difficulties, in particular for apomictic fern species whose spores arise through first division restitution.

The germination trials show that only about 37% of spores in apomictic *C. distans* are able to develop to sporophytes and I find that some spores only germinate to the gametophyte level (although it is possible sporophytes would have arisen had we been able to observe them for longer). Although we might discount these spores as dead ends, observations made during spore growing for cytological work found that gametophytes in *C. distans* will frequently develop runner-like structures that extend a thin line of cells away from the main gametophyte and then develop their own rhizoids and leaves, and may on occasion also give rise to sporophytes (Sosa 2022; pers. obs.); this particular form of vegetative reproduction has not been described before (although it may be referred to in passing by Martínez et al. 2017). Nevertheless, if one of these spores were to arrive in suitable habitat, and given this newly identified form of vegetative reproduction, it would theoretically be able to establish a gametophyte-only population, as has been reported in other fern species (Emigh and Farrar 1977; Pinson et al. 2017; Farrar 1967).

Of particular note—especially given the results of the range analyses—is the finding that larger spores tend to have higher germination and development to sporophyte. Although the predictive power of this relationship is relatively small, these findings point to the importance of spore provisioning. They also align with our finding that individuals with medium-sized spores are the most successful at attaining large ranges, as spores from these plants may be large enough to more often germinate and establish sporophytes, while also being small enough to be transported considerable distances by wind. This is, to my knowledge, the first time that the relationship between fern spore size and germination has been studied, and that the importance of spore provisioning has been highlighted in ferns. Especially given the variation in spore form, the heritability of spore size and of spore forms remains to be explored in further detail to uncover how strongly selection may be able to act on a process that seems so prone to mistakes. Further, I was unable to assess the relationship between mean spore size for the different spore forms because of the way I had coded my data. This remains an interesting question worth exploring, since the different spore forms may not be contributing equally to the variation we observe in size (for example, some spore forms may be generating viable spores with higher size variance than other forms, or may be disproportionately generating smaller or larger spores). It is also worth exploring the hypothesis that, even if selection can only act weakly on sporogenesis—such that variation in spore form remains commonplace—variation in spores as a result of first division restitution may be advantageous enough to be maintained despite its apparent randomness and the costs of lost spores.

## 5. Conclusion

This study was driven by the desire to explore the ever-present question: how do trade-offs shape the variation we observe in nature? Through careful study of spore variation in *C. distans*, I have uncovered yet another example in support of and slightly extending Baker’s rule, as the apomictic lineages of *C. distans* were distributed far more widely than the sexual counterparts. I also found that spore size variation is akin to that observed in pollen size, if not greater—something which has not been documented for ferns before. Of greatest note, however, is the role of this variation. I find that larger spores are both more likely to germinate and develop into sporophytes. Given that smaller spores are expected to travel over longer distances, these two trends establish a clear trade-off with opposing forces that ultimately lead individuals with medium-sized spores to be those with the largest ranges.

The variation in spore size is further enhanced by the modified sporogenesis observed in this species. I find high prevalence of mistakes in *C. distans* sporogenesis, along with generally low germination and establishment rates. It appears these mistakes occur mostly randomly, although the mistakes take on recurrent patterns that I have catalogued into spore forms. Importantly, however, spores generated via aneuploidy can still be viable, and may be an important additional driver generating variation in spore size. The heritability and selection on spore size and spore forms remains to be explored, as does the role of environmental constraints on survival of different groups.

These findings are exciting for several reasons. Not only do we observe quite clear trade-off dynamics between dispersal and offspring survival that appear to affect spore size, but the central role of spore size variation and its drivers are underscored. In light of the fact that many spores that were historically considered abortive are fully viable and likely shaping evolution in important ways, it is worth remarking on what these results illustrate more broadly. What must be highlighted here is that what we considered ‘abortive’, ‘incorrect’, or ‘disabled’ is constructed (Hughes 1999). This does not mean to erase the very real existence of disabled human and non-human relatives, but rather I mean to remark that society constructs what is possible for those who are disabled—what can and cannot even ***be***. Disability in Western society has been historically constructed as a form that is lesser than, ripe to be dominated over (Abberley 1987), and thus something that demands to be cured if we hope to not exist at the bottom of the pecking order (Michalko 2009). Beyond the crushing systemic oppression faced by disabled people, if we also consider the deeply entrenched politics of shame that surround disability in particular (Caron 2008), it becomes essential that we face these preconceptions straight on and that we reconsider what it *means* to be disabled in our society. We have constructed disability in a binary: to be disabled is bad, shameful, wrong; to be abled is good, healthy, prideful. Yet the findings described herein once again show us how narrow and harmful this view is. The way in which we have constructed disability is ultimately affecting how we perceive so-called ‘genetic errors’—both in humans and in other species—thus severely limiting what we even allow ourselves to imagine ‘disabled’ beings may be capable of, and in the case of humans also limiting what disabled people are allowed to become. This matter is especially pressing considering the foundations of evolutionary biology itself, which are steeped in eugenics and ableism (see for example Fisher 1921; or Huxley 1936; see Saini 2019 for an extensive history on this topic; and Powell 2020 for a focus on ableism in particular). Thus, this issue is not one that is *separate from* biological study—it is in fact intrinsically intertwined, and as such it becomes morally imperative that we name these biases and work to correct them in our own research, and in our lives.

## Acknowledgements

First and foremost, apologies and amends are owed to the Traditional Custodians and Indigenous Nations from whose lands samples in this study were taken without consent nor compensation. While I have done my best to be respectful of their non-human relatives (i.e. the ferns studied herein), I have certainly fallen short.

This research would not have been possible without generous funding from the following institutions and organisations: the American Society of Plant Taxonomists, the Biology Department at Duke University, the Botanical Society of America, the International Association for Plant Taxonomy, the National Geographic Society, Sigma Xi, and the Society of Systematic Biologists. Fieldwork was possible thanks to the National Geographic Society’s Young Explorers Grant. I am additionally indebted to all the herbaria who graciously lent me their specimens: State Herbarium of South Australia (AD), Auckland War Memorial Museum (AK), Queensland Herbarium (BRI), Australian National Herbarium (CANB), Manaaki Whenua – Landcare Research (CHR), Australian Tropical Herbarium (CNS), Tasmanian Museum and Art Gallery (HO), Royal Botanic Gardens Victoria (MEL), University of New England (NE), Royal Botanic Gardens and Domain Trust (NSW), Western Australian Herbarium (PERTH), and Smithsonian Institution (US).

I am incredibly grateful to Mark Rausher for welcoming me into his lab and for his steadfast support. His insightful thoughts, along with those of Anne Yoder, Louise Roth, and Paul Manos, significantly improved this work. I owe much gratitude to innumerable colleagues and friends for asking astute questions and making keen observations that have much improved my science; special thanks are due to Ray Allen, Lauren Carley, Jen Coughlan, Tzu-Tong Kao, Devin McCarthy, and Kathryn Picard. An immense debt is owed to Layne Huiet for her assistance with the loans involved in this study, to Ashley Field for his assistance with fieldwork, to Gordon Burleigh for his support with sequencing, and to Julia Notar for her help in reviewing and formatting this paper for publication.

Some of the ideas in this study were first developed under tutelage of Kathleen Pryer and Michael Windham.

## Footnotes

1 Note that I explicitly use the term “range expansion” rather than the more common term “colonisation”. Scientific research and its findings have often been used to claim manifest destiny and justify human actions by settler-colonisers and oppressors (for an analysis of this, see Cardozo and Subramaniam, 2013). In rejecting language laden with historical connotation and legacies of trauma, I hope to underscore that scientific research cannot be used uncritically to justify human actions.

2 Especially because these mistake spores have historically been dismissed, and if we consider the politics of ableism—which is rampant in Western thought and society through its rejection on the ‘variation of being’ (Wolbring 2008) and whose modern conception is strongly rooted in early evolutionary biology and eugenics (Powell 2020)—it is essential that we, as researchers and humans, not dismiss any variation that we do not understand. For these reasons I refer to irregular spores as ‘forms’ rather than errors.

