## Supplemental Information for "The importance of mistakes: Variation in spores reveals trade-offs for range expansion in the xeric-adapted Australasian species *Cheilanthes distans* (Pteridaceae)"

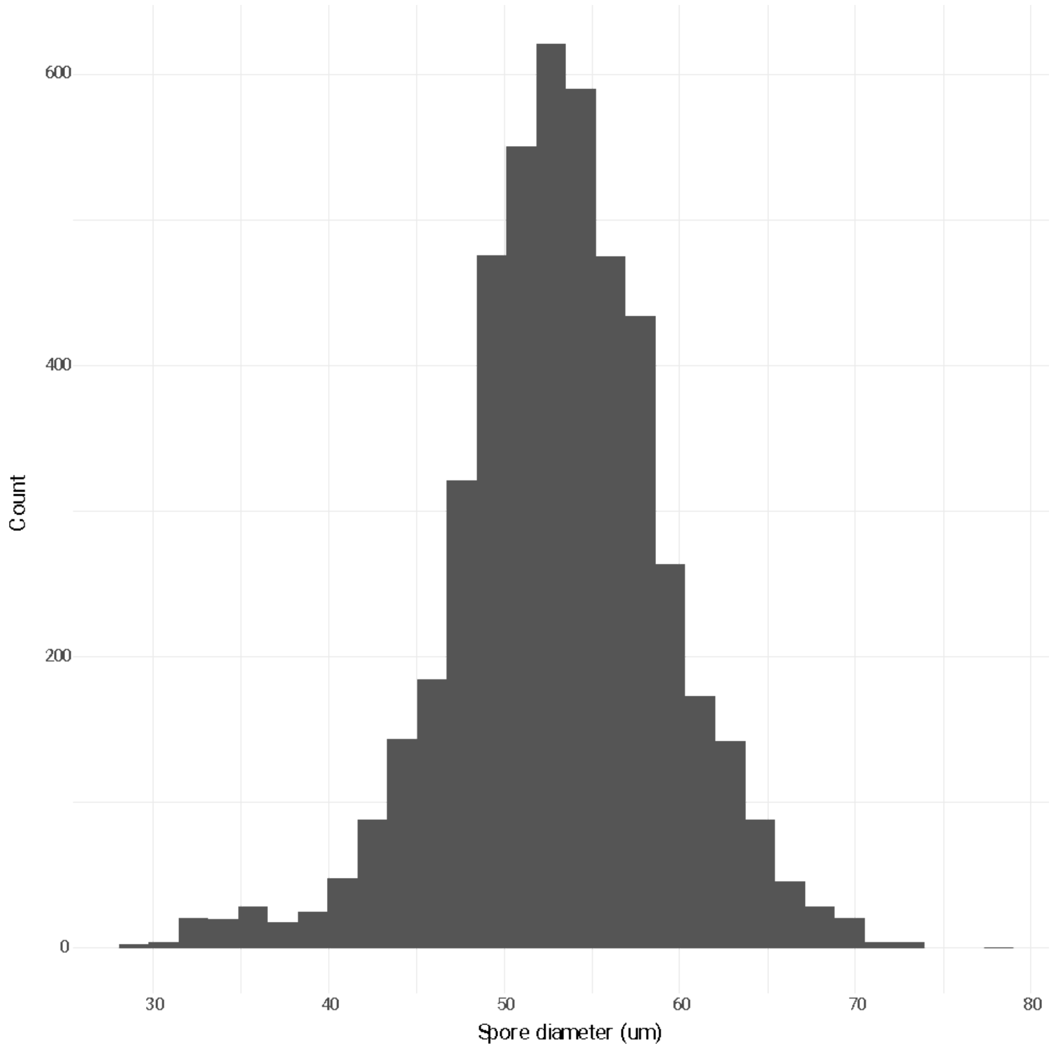


**Figure SI.1: Histogram of all spore diameter measurements for *Cheilanthes distans*.**

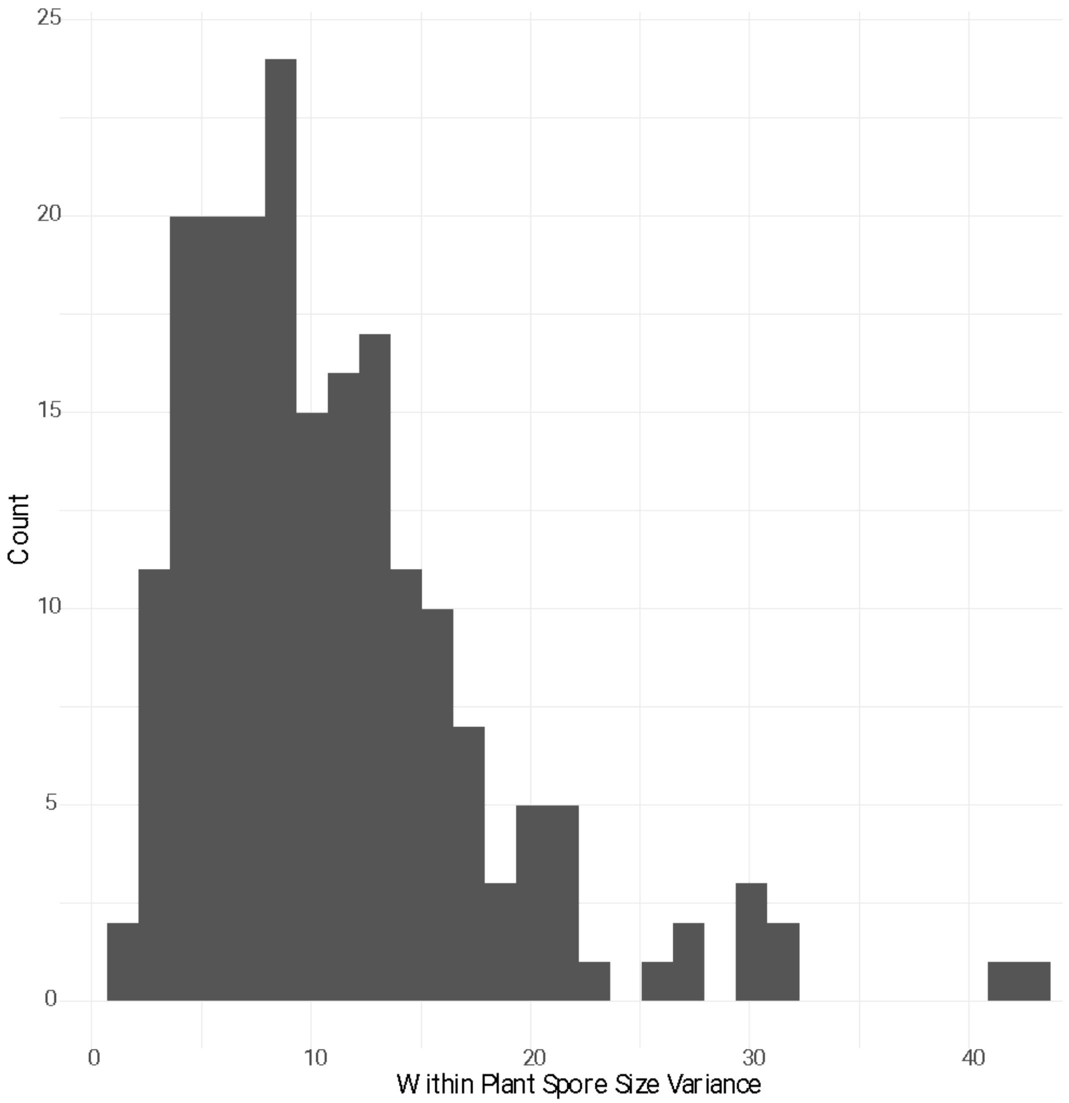


**Figure SI.2: Histogram of the within plant (i.e. within sample) variances in spore diameter measurements for *Cheilanthes distans*.**

**Table SI.1: Correlations between spore forms within a plant, including their confidence intervals.**

| Spore Form | A | B | C | D | E |
|---|---|---|---|---|---|
| A | | | | | |
| B | 0.09 ± 0.80 | | | | |
| C | -0.08 ± 0.81 | 0.01 ± 0.81 | | | |
| D | -0.1 ± 0.80 | -0.04 ± 0.81 | -0.12 ± 0.80 | | |
| E | -0.14 ± 0.79 | -0.18 ± 0.77 | -0.14 ± 0.79 | -0.12 ± 0.80 | |
| F | -0.18 ± 0.78 | -0.34 ± 0.68 | -0.23 ± 0.76 | -0.11 ± 0.80 | 0 ± 0.81 |

**Table SI.2: Summary statistics for growth by spore forms.**

| Form Code | Count | Mean Proportion of Germination | Proportion of Germination SD | Mean Proportion to Sporophyte | Proportion to Sporophyte SD |
|---|---|---|---|---|---|
| B | 4 | 0 | 0 | 0 | 0 |
| C | 5 | 0.130 | 0.181 | 0.130 | 0.181 |
| D | 15 | 0.317 | 0.351 | 0.310 | 0.356 |
| E | 14 | 0.378 | 0.434 | 0.298 | 0.400 |
| F | 19 | 0.625 | 0.342 | 0.604 | 0.341 |
